# Principles of ecDNA random inheritance drive rapid genome change and therapy resistance in human cancers

**DOI:** 10.1101/2021.06.11.447968

**Authors:** Joshua T. Lange, Celine Y. Chen, Yuriy Pichugin, Liangqi Xie, Jun Tang, King L. Hung, Kathryn E. Yost, Quanming Shi, Marcella L. Erb, Utkrisht Rajkumar, Sihan Wu, Charles Swanton, Zhe Liu, Weini Huang, Howard Y. Chang, Vineet Bafna, Anton G. Henssen, Benjamin Werner, Paul S. Mischel

## Abstract

The foundational principles of Darwinian evolution are variation, selection, and identity by descent. Oncogene amplification on extrachromosomal DNA (ecDNA) is a common event, driving aggressive tumour growth, drug resistance, and shorter survival in patients^1-4^. Currently, the impact of non-chromosomal oncogene inheritance—random identity by descent—is not well understood. Neither is the impact of ecDNA on variation and selection. Here, integrating mathematical modeling, unbiased image analysis, CRISPR-based ecDNA tagging, and live-cell imaging, we identify a set of basic “rules” for how random ecDNA inheritance drives oncogene copy number and distribution, resulting in extensive intratumoural ecDNA copy number heterogeneity and rapid adaptation to metabolic stress and targeted cancer treatment. Observed ecDNAs obligatorily benefit host cell survival or growth and can change within a single cell cycle. In studies ranging from well-curated, patient-derived cancer cell cultures to clinical tumour samples from patients with glioblastoma and neuroblastoma treated with oncogene-targeted drugs, we show how these ecDNA inheritance “rules” can predict, *a priori*, some of the aggressive features of ecDNA-containing cancers. These properties are entailed by their ability to rapidly change their genomes in a way that is not possible for cancers driven by chromosomal oncogene amplification. These results shed new light on how the non-chromosomal random inheritance pattern of ecDNA underlies poor outcomes for cancer patients.

---

Inheritance, variation, and selection are foundational principles of Darwinian organismal evolution that have been used to explain how cancers evolve^5-8^. The concept of genetic identity by descent is central to the application of evolutionary theory to cancer, suggesting a physical basis for identity through chromosomal inheritance during cell division – thereby explaining the clonal trajectories commonly seen in tumours^9-12^. However, several issues challenge current models of tumour clonal evolution. First, some aggressive forms of cancer maintain high levels of intratumoural copy number heterogeneity instead of undergoing selective sweeps, as would be predicted^13^. This is especially true for amplified oncogenes, whose cell-to-cell variability remains high, despite the fitness advantage conferred^2,14-16^. Consequently, the mechanisms maintaining heterogeneous oncogene amplification events have been difficult to establish. Second, the ability of some cancers to rapidly adapt to changing conditions, including treatment, by changing their genomes, especially changing the copy number of amplified oncogenes, isn’t well explained by current models of genetic inheritance^2^. Third, the lag time to resistance predicted by the selection for drug resistance-conferring mutations arising in a single cell, or a small number of cells, isn’t seen in some cancers, raising questions about whether tumours are undergoing a genetic bottleneck^2,17^. The presence of extrachromosomal DNA (ecDNA) amplification may explain some of these paradoxical features. Extrachromosomal oncogene amplification on circular particles that lack centromeres is now recognized to be a common event in human cancer that is linked to poor outcome and treatment resistance in patients^1,3^. It has been suggested that ecDNAs, because they lack centromeres, are unequally segregated to daughter cells during cell division^18,19^. However, the impact of non-chromosomal oncogene inheritance in cancer—random identity by descent—on intratumoural genetic heterogeneity, accelerated tumour evolution, enhanced ability to withstand environmental stresses, and rapid genome change on therapeutic resistance, is not well understood. Here, we apply a powerful, integrated tool kit, including mathematical modeling, evolutionary theory, unbiased image analysis, CRISPR-based ecDNA tagging with live cell imaging, and longitudinal analyses of patients’ tumours, to deduce the “rules” of ecDNA inheritance and to reveal the functional consequences.

Chromosomal segregation during mitotic cell division ensures that each daughter cell has the same DNA content (red line, Fig. 1a). If ecDNA segregation is random, then we predict a Binomial (approximately Gaussian) distribution in the per-cell content of ecDNA, post-mitotic division (Fig. 1a, Supplementary Information 1.1). Therefore, we developed a method of using unbiased image analysis to quantify ecDNA in daughter cells after cell division, using FISH probes to detect the amplified oncogenes residing on those ecDNAs, and Aurora B Kinase immunostaining to identify the daughter cells post-mitosis^20^ (Fig. 1b). In cancer cell lines of different histological types, including prostate, gastric, colon cancer cells, and glioblastoma cells, carrying different oncogenes on ecDNA, we quantified the ecDNA distribution of approximately 200 post-mitotic daughter cells per cell line, which permits sufficient resolution (Supplementary Information 1.3), revealing a Gaussian distribution that was independent of cancer cell type or the oncogene contained on the ecDNA (Fig. 1b,c). The fraction of segregated ecDNA per daughter cell (histograms) was highly concordant with the theoretical prediction of random segregation (dashed line) (Kolmogorov-Smirnov (KS) test p > 0.05) (Fig. 1c, Supplementary Information 1.2, 1.3). In one of the cancer cell line models, SNU16, *MYC* and *FGFR2* are found on separate ecDNAs, revealing that oncogenes on different ecDNA segregated independently and randomly (Fig. 1c), adding an additional layer of genetic diversity to tumour cells.

**Figure 1.**
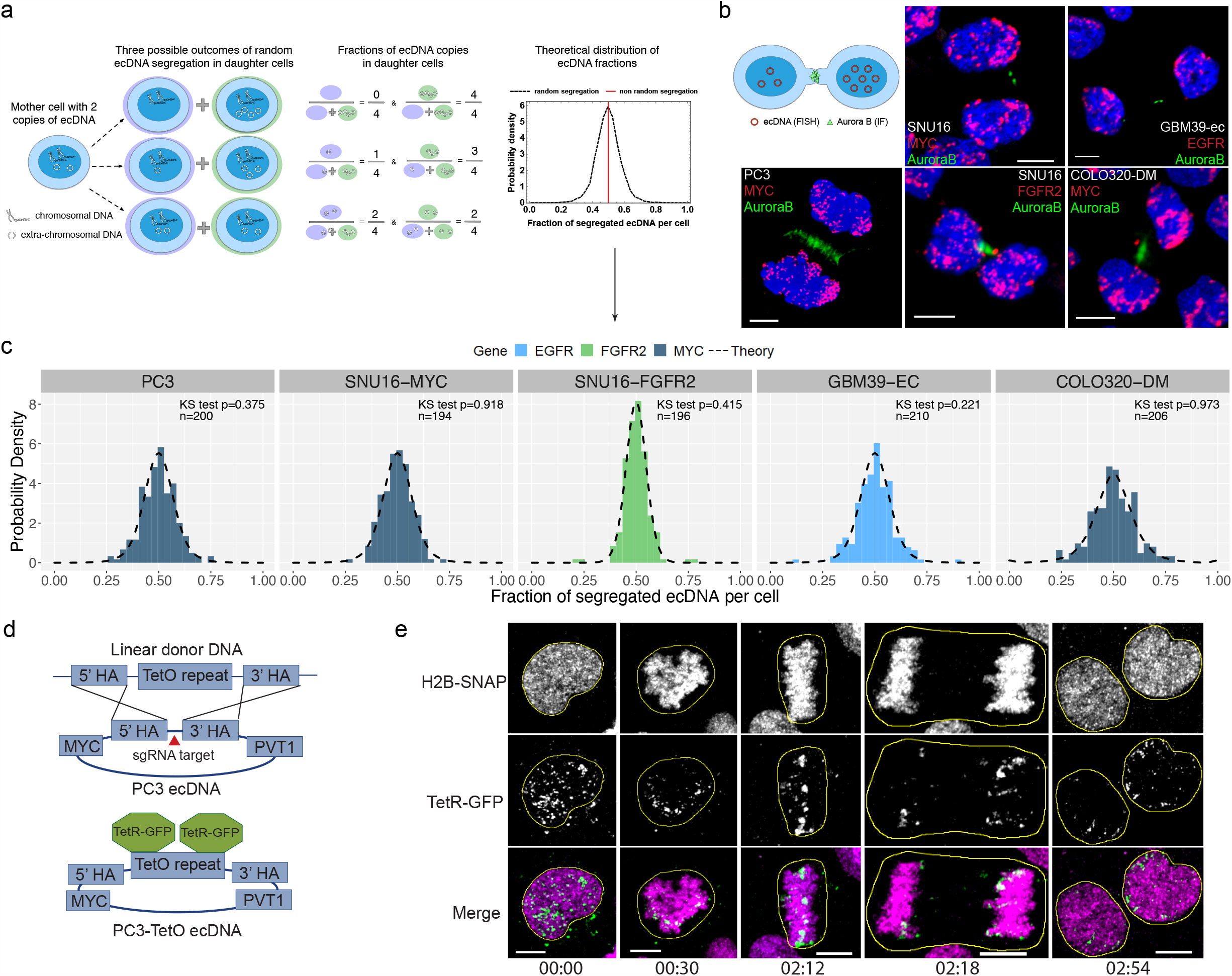
ecDNA is randomly segregated to daughter cells. **a**, Schematic of ecDNA segregation and predicted distribution of ecDNA fractions. **b**, Representative images of ecDNA distribution to daughter cells, identified by Aurora B midbody staining, in multiple cancer cell lines in late-stage mitosis. **c**, Distribution of ecDNA fractions in cancer cell lines analyzed in **b**, showing agreement between theoretical prediction (dashed lines) and observation (histograms) (KS test p values >0.05). **d**, Schematic of CRISPR-based genetic approach used for live-cell imaging of ecDNA in prostate cancer cells. **e**, Live-cell time lapse imaging reveals unequal distribution of ecDNA between daughter cells. Time stamps hh:mm. All scale bars 5µm.

To confirm these correlative observations, we designed a live-cell imaging system to visualize ecDNA dynamics during cell division. We used CRISPR-Cas9^21^ to insert a TetO array into the intergenic region between *MYC* and *PVT1* of the ecDNA in PC3 prostate cancer cells (Fig. 1d). Insertion of this array was confirmed by PCR, sanger sequencing, and TetO-MYC dual FISH (Extended Data Fig. 2a-d). Subsequent expression of TetR-GFP, which binds the TetO array enabled tracking of ecDNA throughout the cell cycle (Fig. 1d). Chromatin was detected by a histone H2B-SNAP tag fusion labeled with the newly developed JF_669_ SNAP tag ligand^22^. Live-cell time-lapse imaging of PC3-TetO cells revealed the random inheritance pattern of ecDNA during cell division (Fig. 1e, Supplementary Video 1).

Having demonstrated that ecDNA drives random identity by descent through random segregation during cell division, we turned our attention to the other pillars of Darwinian evolution – variance and selection. Intratumoural heterogeneity plays a significant role in therapy resistance and tumour evolution^23,24^. To better understand the impact of ecDNA on heterogeneity, we generated a theoretical model of the per-cell distribution of ecDNA (Fig 2a, Supplementary Information 2.1), based on the observed pattern of random segregation. Specifically, starting with a single cell with a single ecDNA, let *N*_*k*_ (*t*) denote the number of cells with *k* ecDNA at time *t*. Assuming independent replication and random segregation, the dynamics of *N*_*k*_ (*t*) are governed by a set of coupled differential equations.

**Figure 2.**
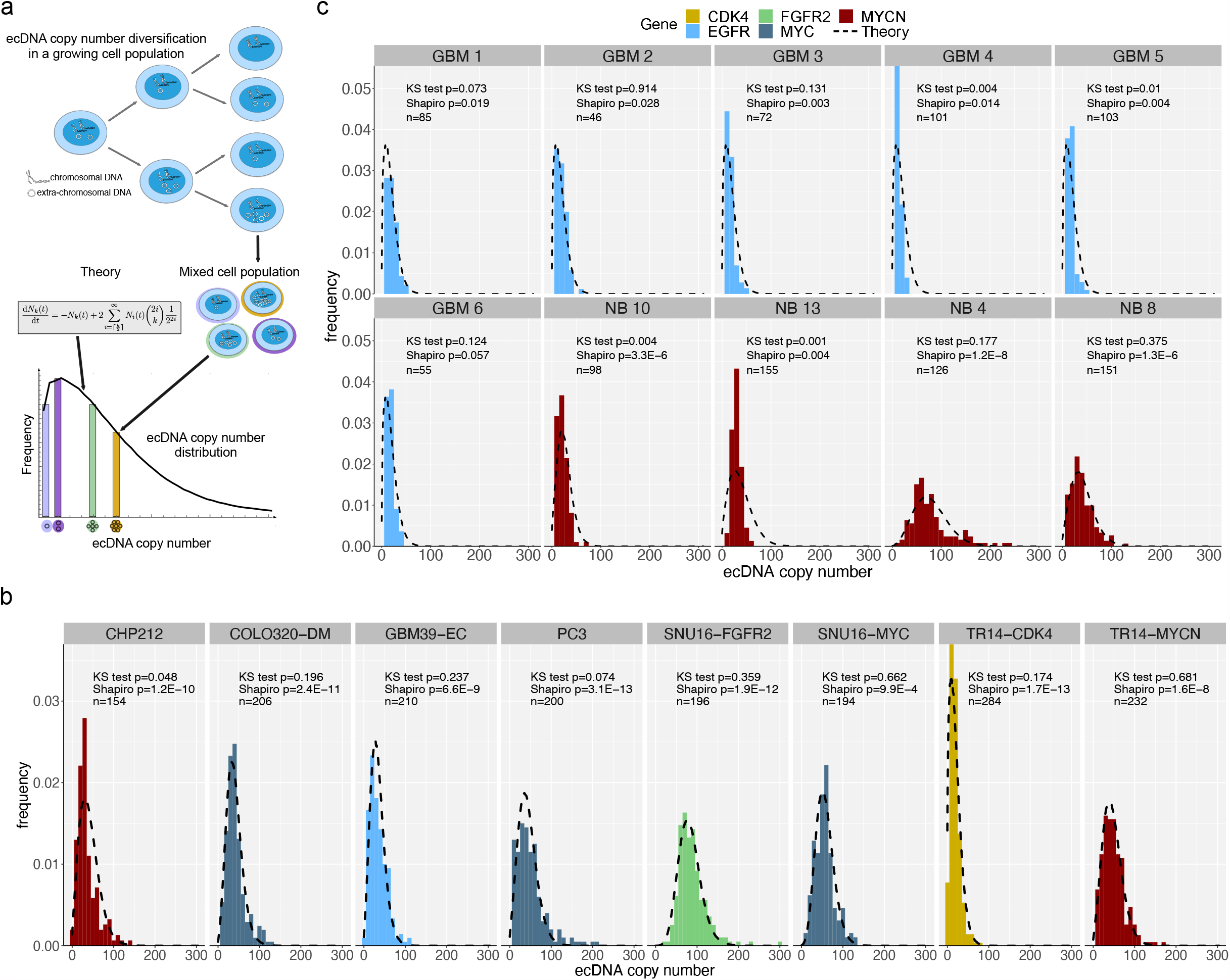
Random segregation of ecDNA promotes intratumoral heterogeneity of oncogenes in cancer cell lines and patient tumor samples. **a**, Schematic showing predicted impact of random ecDNA segregation on single cell oncogene copy number distribution. **b**, Distribution of ecDNA oncogene copy number, assessed by interphase FISH, in cancer cell lines. Agreement between theoretical prediction (dashed lines) and observation (histograms), reveals that oncogene copy number largely follows the prediction distribution (KS test p > 0.05). Shapiro-Wilk p < 0.05 suggests ecDNA number does not resemble a normal distribution. **c**, Distribution of ecDNA oncogene copy number, assessed by interphase FISH, in glioblastoma (GBM) and neuroblastoma (NB) patient tumor samples.

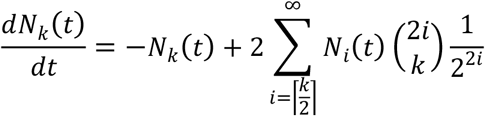

The differential equations can be used to analytically estimate the distribution of ecDNA numbers per cell in a growing tumour population. To test the dynamics, we quantified ecDNA copy number distributions from 6 ecDNA^+^ lines of different cancer types, bearing different amplified oncogenes on ecDNA — two lines contain two distinct species of ecDNA as indicated (Fig. 2b). We observed a wide distribution of copy number in each cancer cell line, with variation primarily dependent on the mean copy number in each cell line model. The observed ecDNA copy number distributions were clearly non-Normal (Fig. 2b; Shapiro Wilk p < 0.05) and matched the predicted analytical distribution (KS test p > 0.05), except for inflation at extreme values in a few cell-lines (Supplementary Information 1.2). The inflation is likely due to positive selection as described below.

We next sought to test whether ecDNA heterogeneity can be observed and modeled in patient tumour samples. We received FISH images on patient tumour samples or patient tumour tissue from 6 GBM^2^ and 14 neuroblastoma (NB) patients. These tumours were suspected of having ecDNA amplification of either EGFR or MYCN, respectively, due to their extremely high copy number—copy number greater than 16 has been found to be almost exclusively due to ecDNA amplification^1^. We quantified the distribution of ecDNA FISH signals in these patient samples and observed distributions that again showed extreme cell-to-cell variation with a non-Normal distribution (Fig. 2c, Extended Data Fig. 3a), strongly suggestive of positive selection in vivo, but remained in strong agreement with the analytic distributions for most samples (KS test p > 0.05). Small discrepancies can possibly be attributed to underestimation of counts due to the much more limited resolution and number of cells quantified (Supplementary Information 1.3).

Importantly, the significant divergence from a normal distribution (Shapiro Wilk test p < 0.05), is indicative of a power-law tail shift, or overrepresentation of extremely high copy number cells in line with our predicted modeling of ecDNA (Supplementary Information 1.2). Furthermore, the shift to high copy ecDNA suggests that there may be an important role for selection.

To understand whether there is positive selection for ecDNA and to determine how it shapes tumour evolutionary dynamics, we simulated the expansion of a single cell colony with a single ecDNA into a population of 10^5^ cells (Supplementary Information 1.1). Due to random segregation, cells with low ecDNA copy number frequently give rise to a daughter cell without ecDNA. Under neutral selection, this cell is not disadvantaged. Consequently, ecDNA prevalence rapidly decays to a small minority of cells, consistent with the rare observation of ecDNA in normal cells (Fig. 3a,b, Supplementary Information 4.1)^3^. In conditions where ecDNA is positively selected, however, the simulations show that ecDNA remains frequent, with a continued presence in a vast majority of the cells (Fig. 3a,b). We compared these simulated data to our empirical measurements of ecDNA prevalence in the cell lines and patient samples measured in Figure 2. In all samples, ecDNA prevalence levels suggested strong positive selection for ecDNA (Fig. 3c).

**Figure 3.**
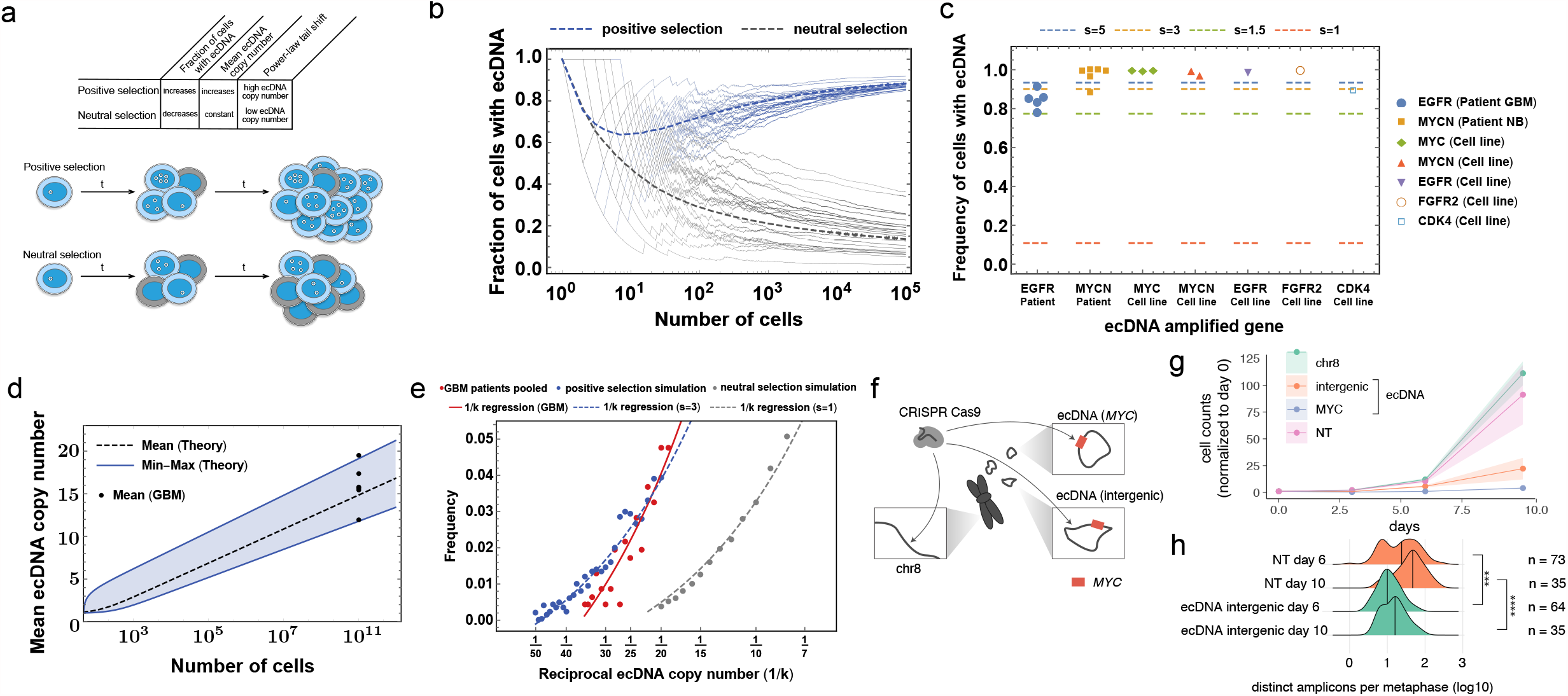
Strong selection for ecDNA in cancer. **a**, Schematic comparing predicted impact of ecDNA under positive or neutral selection. **b**, Simulations showing ecDNA prevalence in populations derived from a single ecDNA+ cell with ecDNA under positive or neutral selection. Positive selection s=3; neutral s=1. **c**, Empirical data comparing the frequency of ecDNA+ cells in cancer cell line and patient samples (colored shapes) to various selection strengths (dashed lines), shows strong evidence for positive ecDNA selection across oncogenes and cancer types. **d**, Comparison between the minimum, maximum and mean copy number predicted by strong ecDNA selection pressure (blue cone) and the ecDNA copy number reached in modeled GBM tumors (blue cone and lines) and the observed mean ecDNA copy number detected in 6 GBM patient samples (dots). **e**, Power law tail shift of ecDNA copy number in GBM patients indicative of strong positive selection. **f**, Depiction of CRISPR-based strategy to test selective advantage given to COLO320DM cells by MYC ecDNA. Arrows indicate regions targeted by sgRNA. **g**, Genome editing of MYC encoded on ecDNA causes massive decrease in cell number that exceeds the impact of intergenic editing, indicative of strong selection for oncogenes on ecDNA. **h**, Quantification of ecDNA numbers per metaphase at 6 and 10 days post CRISPR transfection. P values calculated using Mann-Whitney tests. ***p≤0.0005; ****p≤0.00005.

To better understand the selection landscape of ecDNA, we modeled the predicted ecDNA copy number under strong positive selection, where cells carrying ecDNA are 3 times (s=3) more likely to divide compared to cells with no ecDNA. The simulations predict an exponential increase in the average copy number per cell (Fig. 3d). Remarkably, when plotted against the observed copy number averages in GBM, we again saw strong agreement between the predicted model and our observations (Fig. 3d). Interestingly, these samples fit with the modeled tumour growth when the tumour reaches a size reasonable for clinical detection (10^11^ cells), potentially suggesting ecDNA as an early event in the development of these tumours. An additional prediction of our simulations relates to the power-law tail shift (Supplementary Information 2.3,3.1), or overrepresentation of extremely high copy number cells, predicted in ecDNA^+^ populations (Supplementary Information 4.1). We modeled this feature by plotting the distribution of reciprocal ecDNA copy number in simulated populations under either positive selection or neutral evolution (Fig. 3e). When we overlaid the simulated distributions with data from the GBM patient samples, we saw a strong left-shift indicative of strong positive selection (Fig. 3e).

To complement the evolutionary analyses showing ecDNA selection, we designed a set of CRISPR studies to determine the reliance of tumours on ecDNA and on the oncogenes encoded within the ecDNAs for growth. We designed sgRNAs targeting different genomic regions of COLO320-DM *MYC* ecDNA (intergenic region on ecDNA and *MYC* gene body on ecDNA) and a non-amplified, intergenic region of chromosome 8 (Fig. 3f). We infected the cells with Cas9 and the sgRNAs by lentiviral vectors, quantifying cell proliferation and ecDNA copy number. While Cas9-targeted cutting of chromosome 8 showed minimal impact on cell proliferation, targeting of the ecDNA on an intergenic region, and even more so on *MYC* on the ecDNA, caused an extreme growth deficit (Fig. 3g). When we quantified ecDNA copy number in these cells, we saw a significant decrease in ecDNA 6 days after initial infection (Fig. 3h, Extended Data Fig. 3b). These data together confirm that ecDNAs, and the oncogenes contained therein, are under strong selective pressure, which influences the mean ecDNA oncogene copy number and per cell distribution in tumours.

Having shown that ecDNA contributes to each of the three pillars of Darwinian evolution — inheritance (i.e. random identity by descent), variation, and selection — in a unique fashion relative to chromosomal inheritance, we asked whether these ecDNA features enable more rapid tumour adaptation to stress than possible through chromosomal inheritance (Fig. 4a). We utilized an isogenic cell line pair derived from a GBM patient^2^ to examine the importance of ecDNA in driving rapid adaptation. GBM39-EC is a patient-derived neurosphere model with a mean copy number of approximately 100 copies of *EGFRvIII*, a gain of function *EGFR* mutation residing on ecDNA^3,4^. GBM39-HSR is an isogenic model, in which all the *EGFRvIII* amplicons reside on chromosomal HSRs, at the same mean copy number with the same DNA sequence (Extended Data Fig. 4a)^4^. Importantly, the heterogeneity of *EGFRvIII* copy number in GBM39-EC correlates with the heterogeneity of EGFRvIII protein expression assessed by flow cytometry (Extended Data Fig. 4b,c). GBM39-EC cells are highly glycolytic^2^. Therefore, we tested the differential effect of glucose restriction on GBM39-EC and GBM39-HSR cells. We withdrew 80% of normal glucose levels from the culture medium and saw a striking difference — the GBM39-HSR cells were exquisitely sensitive to glucose withdrawal, whereas the GBM39-EC showed no significant decrease in cell growth (Fig. 4b). This ability of GBM39-EC cells to adapt to glucose restriction was mirrored by a rapid decrease in the mean level and overall distribution of *EGFRvIII*-containing ecDNAs per cell (Fig. 4c). Remarkably, this genomic shift took place within a couple of cell cycles. In contrast, the GBM39-HSR cells, which were highly sensitive to glucose restriction, were not capable of rapidly changing their *EGFRvIII* copy number (Fig. 4c).

**Figure 4.**
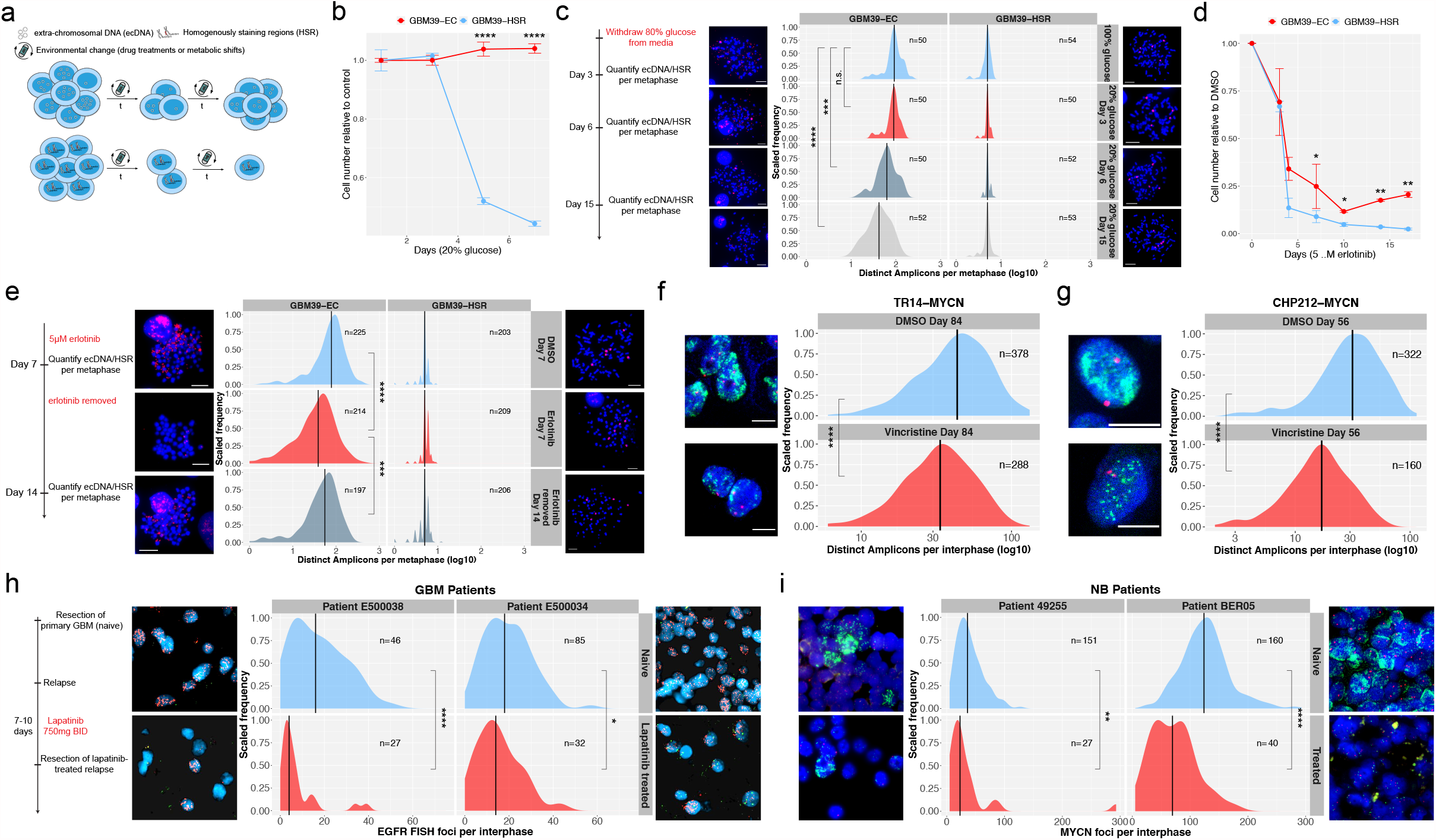
Non-chromosomal inheritance of ecDNA promotes rapid adaptation and resistance to glucose withdrawal and targeted drug treatment. **a**, Schematic depicting how the random segregation of ecDNA and ensuing heterogeneity can drive rapid adaptation and resistance.**b**, ecDNA-containing GBM cells are relatively resistant to glucose withdrawal, whereas GBM cells in which the same oncogene has lodged onto chromosomal loci at near identical copy number (GBM39-HSRs) cannot tolerate glucose withdrawal. Significance determined by two-sided t-tests. Error bars indicate standard deviation. Each data point indicates mean of 3 biological replicates. **c**, Adaptation of ecDNA containing cells to glucose withdrawal is linked to a rapid shift in the distribution of amplicons per cell, unlike the highly sensitive HSR containing cells that cannot modulate amplicon copy number. Timeline of experiment is depicted at the left of the panel. Red FISH signal is from EGFR FISH probe. **d**, GBM cells with GFRvIII amplified on ecDNA, after an initial response, rapidly become resistant to the EGFR tyrosine kinase inhibitor erlotinib, where-as the GBM39-HSR cells remain highly sensitive. Significance determined by two-sided t-tests. Error bars indicate standard deviation. Each data point indicates mean of 2 biological replicates (4 for day 7). **e**, GBM cells with EGFRvIII amplified on ecDNA rapidly shift the distribution of EGFRvIII amplicons per cell, measured at 7 days, which can also be rapidly reversed within one week by drug withdrawal. Timeline of experiment is depicted at the left on the panel. Red signal is EGFR FISH probe. **f**, NB cell line TR14 shift the copy number distribution of MYCN ecDNA when treated with 43nM vincristine for 12 weeks. Green signal is MYCN FISH probe. **g**, NB cell line CHP-212 shift the copy number distribution of MYCN ecDNA when treated with 5.3nM vincristine for 8 weeks. Green signal is MYCN FISH probe. **h**, Comparison of the distribution of EGFR amplification per cell in two GBM patients before therapy (naive) and after 7-10 days of lapatinib treatment. Red FISH signal is from EGFR FISH probe. Green FISH signal is from Chr. 7 control probe. **i**, Comparison of MYCN ecDNA copy numbers assessed by MYCN (green) FISH in two NB patients before and after recieving chemotherpy including Vincristine. Red signal from Chr. 2 control FISH probe. Scale bars represent 5µm in all images. P values calculated using Mann-Whitney tests except where indicated. N.s. not significant; * p ≤ 0.05; ** p ≤ 0.005; *** p ≤ 0.0005; **** p ≤ 0.0005.

We had previously shown that GBM39-EC cells could become reversibly resistant to the EGFR tyrosine kinase inhibitor (TKI) erlotinib, by lowering ecDNA copy number. Therefore, we examined whether GBM39-EC cells would develop resistance to erlotinib more rapidly than GBM39-HSR cells. Similar to glucose deprivation, GBM39-EC adapted to the changing condition by altering its ecDNA copy number. After initially decreasing in cell number, the GBM39-EC cells became resistant to erlotinib after just two weeks of treatment, shifting their per cell ecDNA distribution in a reversible fashion (Fig 4d,e). In contrast, the GBM39-HSR cells did not shift *EGFRvIII* chromosomal copy number and remained highly sensitive to erlotinib (Fig. 4d,e). We then analyzed two samples taken from GBM patient tumours, as previously described^2^. We compared the primary tumour resection (naïve) to the resected relapse which was treated with EGFR TKI lapatinib for 7-10 days prior to resection. We found a significant decrease in mean EGFR copy number and in the ecDNA distribution in these patients’ tumours (Fig. 4h). To extend our analysis to other ecDNA-containing cancer types, we studied the effect of vincristine, a chemotherapeutic that antagonizes *MYCN* amplification^25^. In vitro, neuroblastoma cell lines TR14 and CHP212 with *MYCN* amplified on ecDNA responded to vincristine by left-shifting the ecDNA distribution, (Fig. 4f,g). When we compared treatment-naïve neuroblastoma biopsies with primary tumour resections after treatment including vincristine, we found a similarly significant decrease in the mean copy number and a left-shift in the ecDNA distribution of *MYCN* in both of these patient tumours, in parallel with the cell line data (Fig. 4i). Interestingly, when CHP212 was treated with the CDK4 inhibitors Abemaciclib, and to a greater extent Palbociclib, a right-shift in the distribution of CDK4 ecDNA was detected in resistant tumour cells (Extended Data Fig. 4d,e, Extended Data Fig. 5).

Together, these data indicate a clear pattern in which ecDNA enables high levels of heterogeneity, which enable increased initial resistance to environmental or therapeutic challenges. Further, the ongoing random inheritance of ecDNA-based oncogenes causes rapid adaptation and the formation of resistance, through a mechanism which is impossible in cells driven by chromosomal alterations.

ecDNA has emerged as a major challenge that forces reconsideration of our basic understanding of cancer. Emerging data demonstrate that the altered topology of ecDNA drives enhanced chromatin accessibility and rewires gene regulation to drive oncogenic transcription^4^. Further, the unique higher-level organization of ecDNA particles into hubs^26^ further contributes to ecDNA-mediated pathogenesis. The findings presented here reveal that ecDNA uniquely shapes each of the foundational principles of Darwinian evolution – random inheritance by descent, enhanced variation through random segregation, and selection, thereby accelerating tumour cell evolution to maximize adaptation. Treating such cancers may require targeting the unique adaptability of ecDNAs in the future.

## Acknowledgements

P.S.M. was supported by a grant from The National Brain Tumour Society. Supported by. NIH R35-CA209919 (H.Y.C.). H.Y.C. is an Investigator of the Howard Hughes Medical Institute. BW is supported by a Barts Charity Lectureship (grant MGU045). The UCSD microscopy core is supported by NINDS NS047101.

## Author Contributions

J.T.L., V. B, B.W., and P.S.M. conceived the project. J.T.L., C.Y.C, L.X., J.T., K.L.H., K.E.Y., Q.S., and M.L.E., performed experiments. Y.P., W.H., V.B., and B.W. performed computational modelling. J.T.L., C.Y.C., Y.P., L.X., J.T., K.L.H., K.E.Y., Q.S., and U.R. analysed data, guided by S.W., C.S., Z.L., W.H., H.Y.C., V.B., A.G.H., B.W., and P.S.M. J.T.L., W.H., V.B., B.W., and P.S.M. wrote the manuscript with feedback from all authors.

---

**Extended Data Fig. 1.**
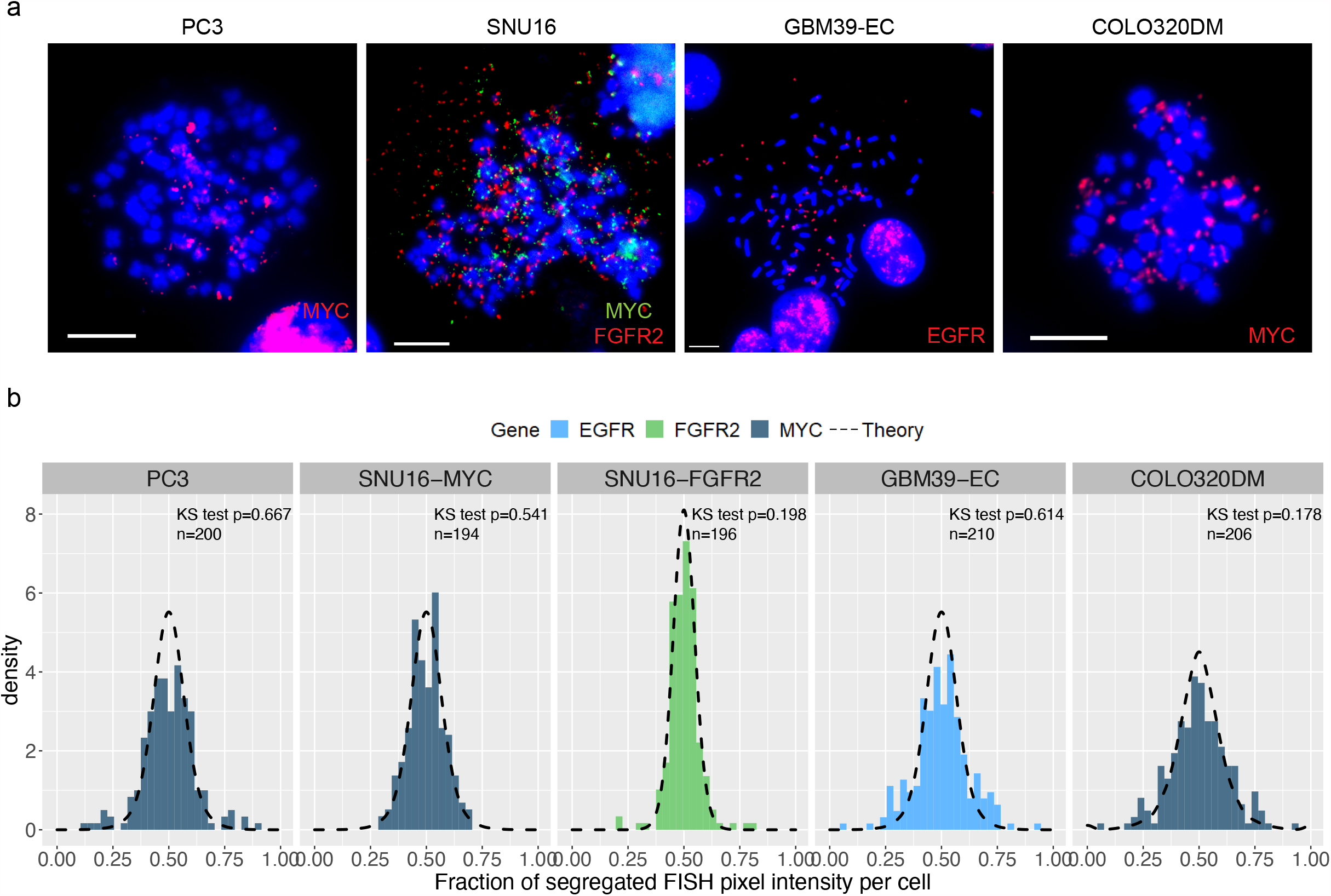
Quantification of ecDNA pixel intensity shows uneven random segregation to daughter cells. **a**, Representative metaphase FISH images for cell lines used to quantify segregation dynamics in **Fig. 1. b**, The same daughter cells analyzed in **Fig.1c** were analyzed by quantifying the pixel intensity of FISH signal in each daughter cell, as a proxy for ecDNA number. Analysis was unbiased and useful for cases in which ecDNA were packed together making counting distinct foci difficult. Agreement between theoretical predictions (dashed lines) and observation (histograms) shown by KS test p value > 0.05. Scale bars 10µm.

**Extended Data Fig. 2.**
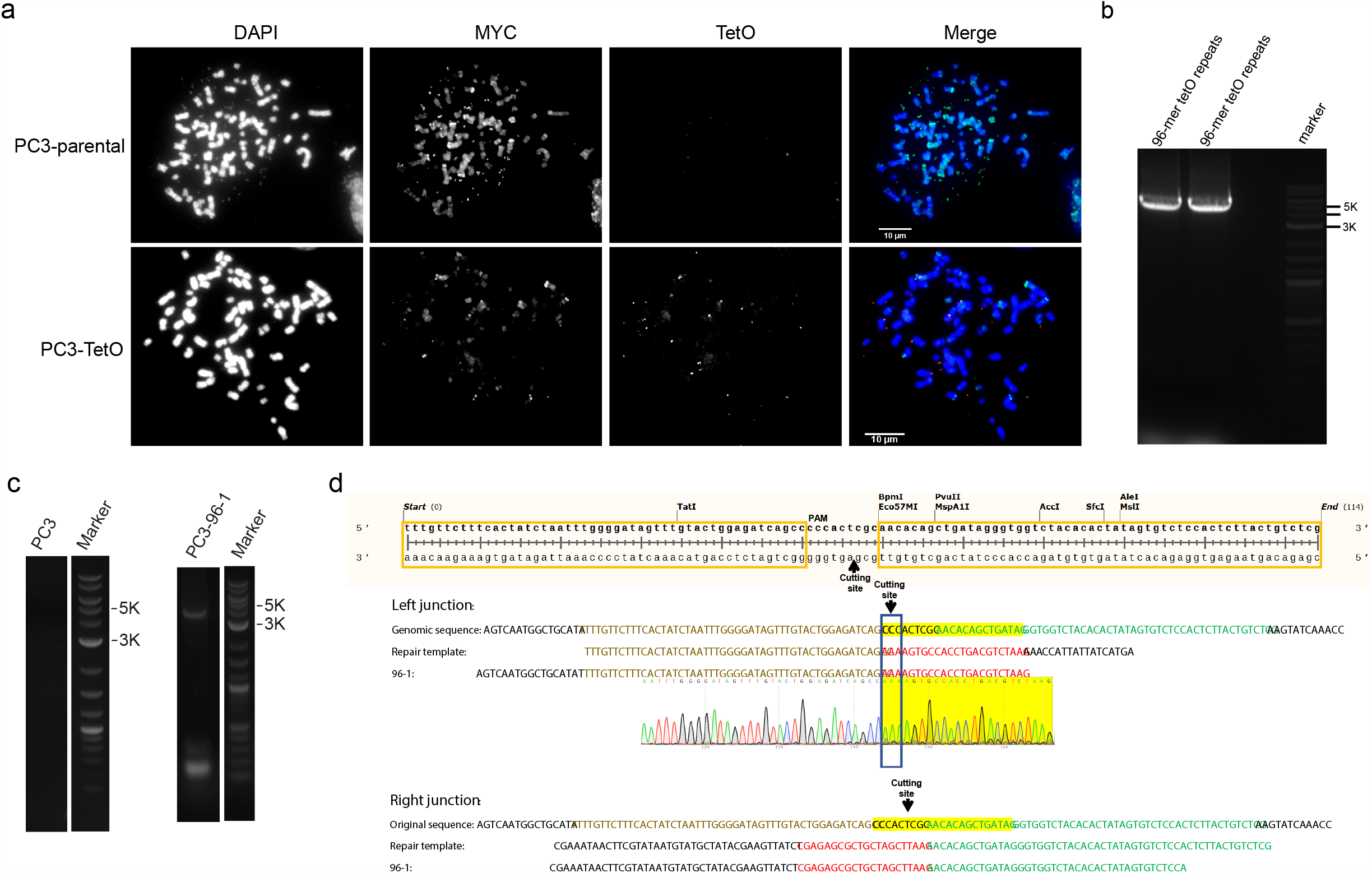
Live cell tracking of ecDNA through insertion of Tet-O array into the ecDNA of PC3 cells. **a**, Representative images of PC3 parental and PC3-TetO cell lines showing extensive MYC amplification on both. PC3-TetO shows significant TetO FISH signal on multiple ecDNA bodies as well. **b**, PCR amplification of 96-mer TetO repeats. **c**, PCR amplification of 96-mer TetO repeats from DNA isolated from PC3-TetO cells confirming insertion. **d**, Sanger sequencing of PCR amplification product from PC3-TetO cells. Both left and right junctions were repaired by homologous recombination at the insertion site.

**Extended Data Fig. 3.**
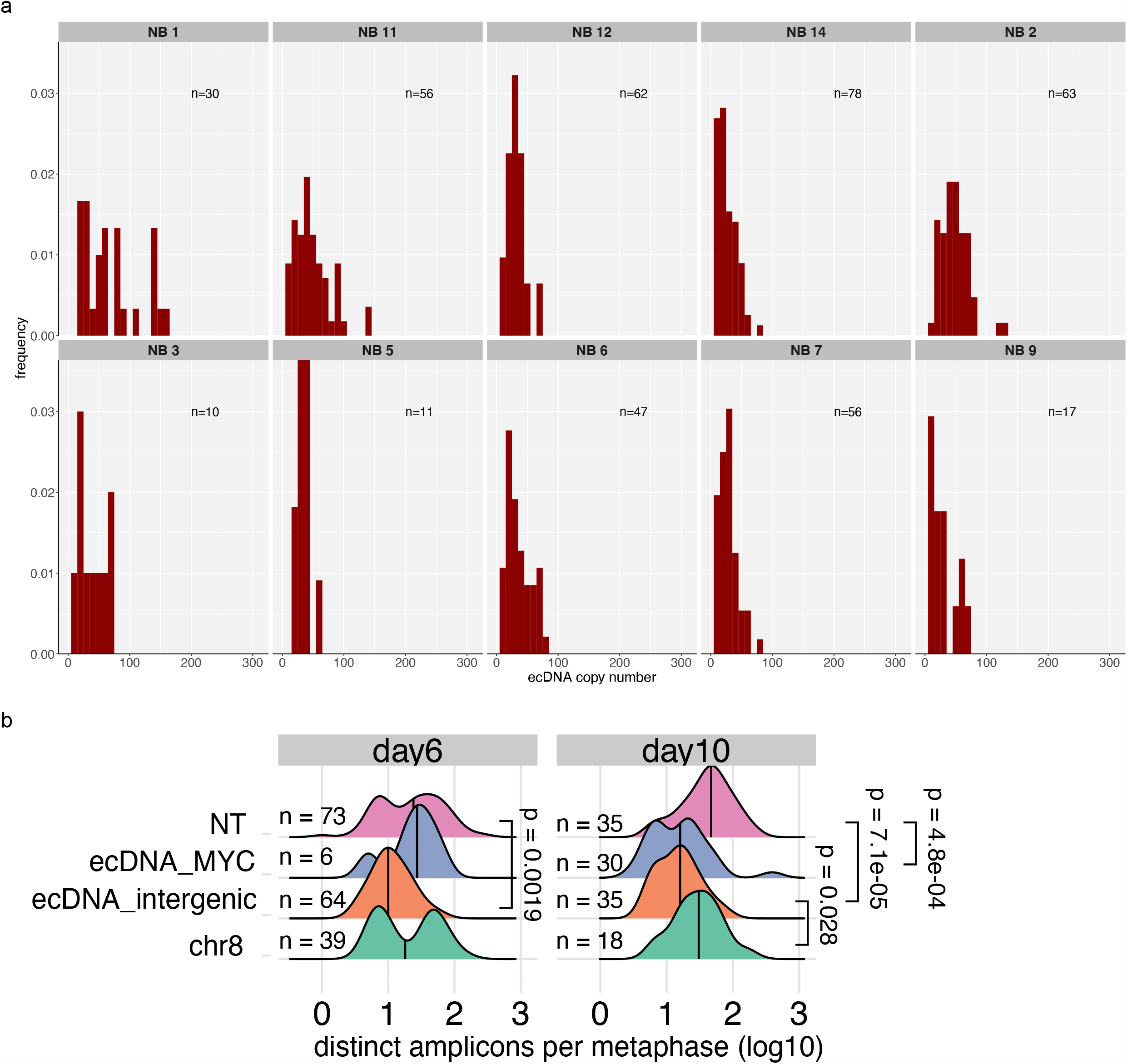
ecDNA heterogeneity and selection. **a**, Histograms of ecDNA copy number assessed by interphase FISH on patient tumor tissue from neuroblastoma (NB) patients. **b**, Quantification of ecDNA numbers at Day 6 and Day 10 after CRISPR cutting of regions of the COLO320-DM genome, either on or off of ecDNA. Shows clear evidence for selection of ecDNA both by the severe drop in copy number when targeted and the inidcation that the copy number begins to return to inital levels. Note ecDNA_MYC at day 6 is severely limited in its growth and only 6 metaphases were able to be identified and imaged.

**Extended Data Fig. 4.**
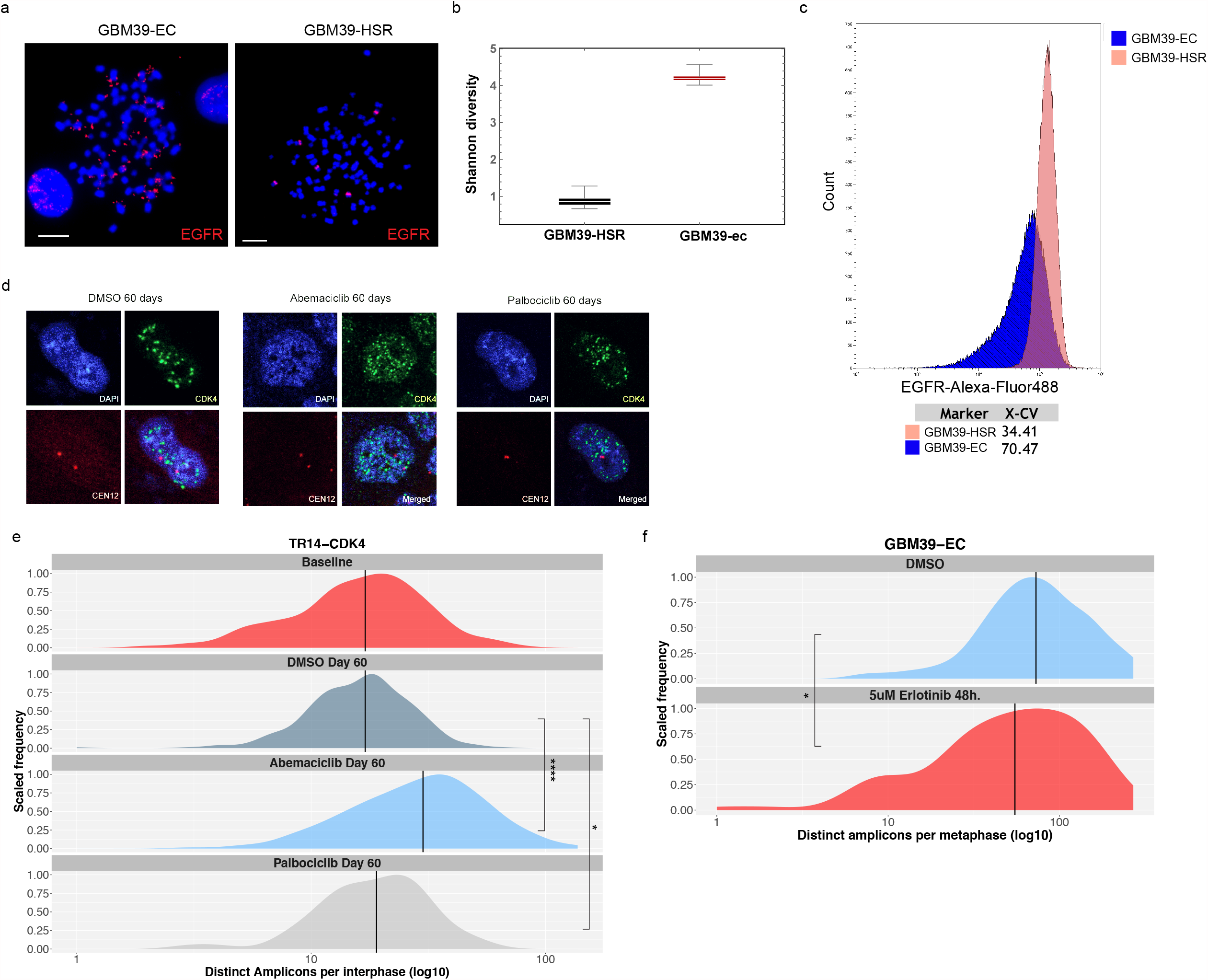
ecDNA dynamically responds to therapeutics. **a**, Representative images of metaphase spread FISH from isogenic GBM39 cell line. **b**, Quantification of the shannon diversity index between isogenic GBM39-HSR GBM39-EC cell lines based on counts of ecDNA amplicons per cell. **c**, Flow cytometry analysis of EGFR protein expression in isogenic GBM39-EC and GBM39-HSR cell lines shows pattern of heterogeneity similar to that seen in copy number. X-CV quantifies the % coefficient of variation for the two samples. **d**, Representative images of TR14 cells treated with Abemaciclib or Palbociclib for 60 days. CDK4 FISH signal shown in green, CEN12 control FISH probe shown in red. **e**, Quantification of experiment described in **c** shows significant shift in CDK4 ecDNA copy number distribution under both drug conditions. **f**, Quantification of EGFR ecDNA in GBM39-EC cells after short-term treatment with erlotinib shows rapid change in ecDNA copy number distribution. Lines indicate medians. P values calculated using Mann-Whitney tests. * p≤0.05; **** p≤0.0001. Scale bars 10 µm.

**Extended Data Fig. 5.**
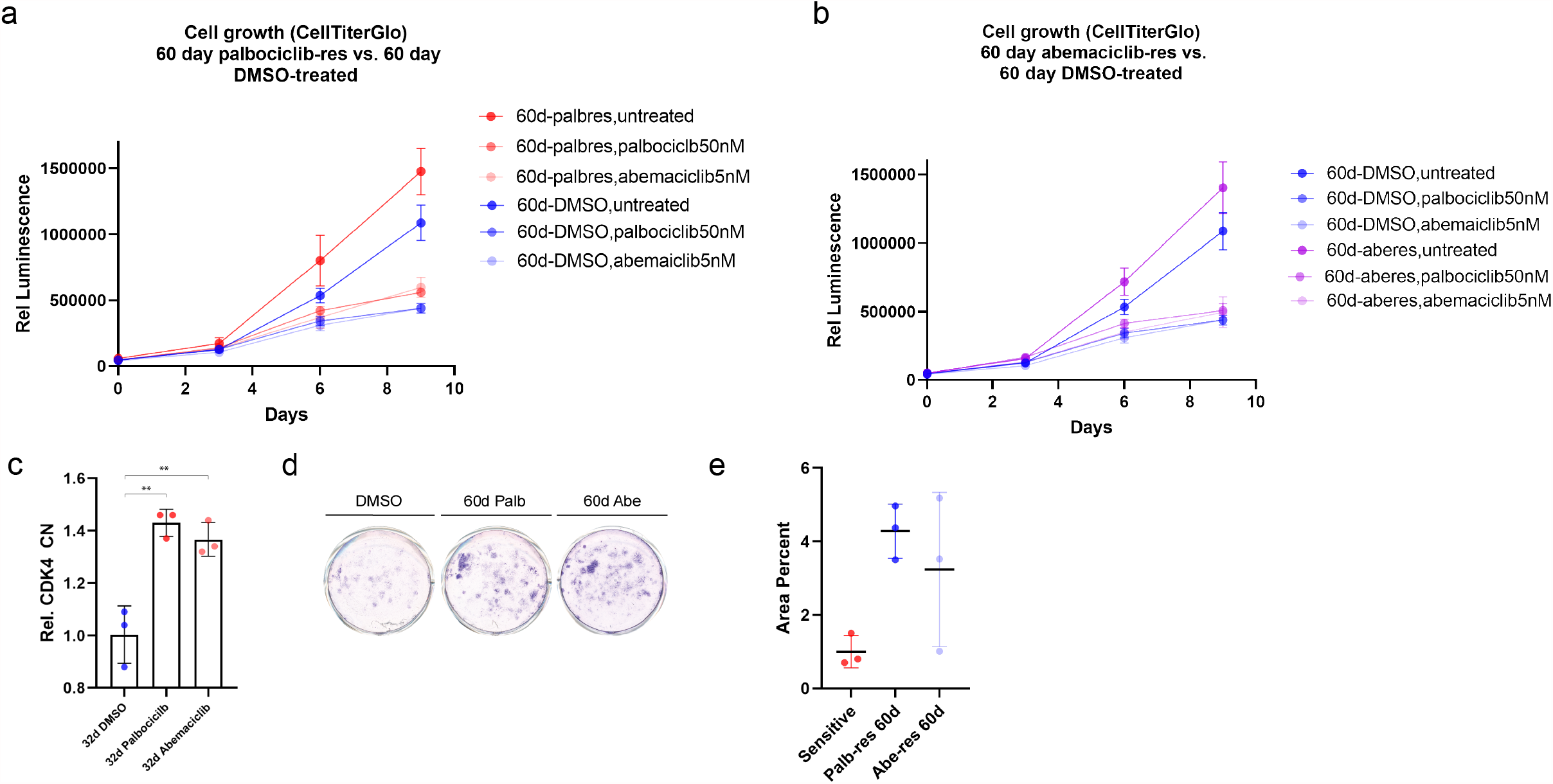
ecDNA dynamics correlate with formation of resistance. **a**, Treatment of long term palbociclib resistant populations of TR14 cells with palbociclib or abemaciclib, showing resistance to treatment. **b**, Treatment of long term abemaciclib resistant populations of TR14 cells with palbociclib or abemaciclib showing resistance to treatment. **c**, Validation of increased ecDNA copy number by qPCR for CDK4. **d**, Crystal violet staining of TR14 cells re-challenged with palbociclib or abemaciclib after development of resistance, or not (DMSO). **e**, Quantification of **d** showing resistance in populations treated with CDK4 inhibitors for 60 days.

**Supplemental Movie 1. Live-cell imaging of PC3-TetO cells**. Live-cell imaging of PC3-TetO cells with chromatin labelled by H2B-SNAP (purple) and ecDNA labelled in green (GFP).

## METHODS

### Cell culture

Cell lines were purchased from ATCC or DSMZ-German Collection of Microorganisms and Cell Cultures (Leibniz Institute) or were a kind gift from J.H. Schulte. GBM39-HSR and GBM39-EC were derived from a patient GBM as previously described (Nathanson cite).

PC3 cells were cultured in DMEM with 10% fetal bovine serum (FBS). COLO320-HSR and COLO320-DM were cultured in DMEM/F12 50%:50% with 10% FBS. SNU16 were grown in RPMI-1640 with 10% FBS. GBM39-HSR and GBM39-EC neurospheres were grown in DMEM/F12 with B27, Glutamax, Heparin (5μg/ml), EGF (20ng/ml), and FGF (20ng/ml). TR-14 cells were grown in RPMI-1640 with 20% FCS. TR-14 cells were cultured in RPMI-1640 with 20% FCS. Cell numbers were counted with a TC20 automated cell counter (Bio-Rad). For drug treatments, drug was replaced every 3-4 days.

### Metaphase chromosome spreads

Cells were concentrated in metaphase by treatment with KaryoMAX colcemid (Gibco) at 100ng/ml for between 3 hours and overnight (depending on cell cycle speed). Cells were washed once with PBS and a single cell suspension was incubated in 75mM KCl for 15 minutes at 37°C. Cells were then fixed with Carnoy’s fixative (3:1 methanol:glacial acetic acid) and spun down. Cells were washed with fixative 3 additional times. Cells were then dropped onto humidified glass slides.

### Fluorescence *in situ* hybridization (FISH)

Fixed samples on coverslips or slides were equilibrated briefly in 2x SSC buffer. They were then dehydrated in ascending ethanol concentrations of 70%, 85%, and 100% for approximately 2 minutes each. FISH probes were diluted in hybridization buffer (Empire Genomics) and added to the sample with addition of a coverslip or slide. Samples were denatured at 72°C for 2 minutes and then hybridized at 37°C overnight in a humid and dark chamber. Samples were then washed with 0.4x SSC then 2x SSC 0.1% Tween-20 (all washes approximately 2 minutes). DAPI (100ng/ml) was applied to samples for 10 minutes. Samples were then washed again with 2x SSC 0.1% Tween-20 then 2x SSC. Samples were briefly washed in ddH_2_O and mounted with Prolong Gold. Slides were sealed with nail polish.

### Dual immunofluorescence – fluorescence *in situ* hybridization (IF-FISH)

Asynchronous cells were grown on poly-l-lysine coated coverslips (laminin, for GBM39-EC). Cells were washed once with PBS and fixed with cold 4% paraformaldehyde (PFA) at room temperature for 10-15 minutes. Samples were permeabilized with 0.5% Triton-X in PBS for 10 minutes at room temperature and then washed with PBS. Samples were then blocked with 3% BSA in PBS-0.05% Triton-X for 30 minutes at room temperature. Samples were incubated in primary antibody, diluted in blocking buffer, for either 1 hour at room temperature or overnight at 4°C. Samples were washed thrice in PBS-0.05% Triton-X. Samples were incubated in secondary antibody, diluted in blocking buffer, for 1 hour at room temperature (all subsequent steps in the dark) and then washed thrice in PBS-0.05% Triton-X. Cells were washed once with PBS and re-fixed with cold 4% PFA for 20 minutes at room temperature. Cells were washed once with PBS then once with 2x SSC buffer. FISH proceeded as described above with the following difference: denaturation was performed at 80°C for 20 minutes.

### Microscopy

Conventional fluorescence microscopy was performed using an Olympus BX43 microscope; images were acquired with a QI-Click cooled camera. Confocal microscopy was performed using a Leica SP8 microscope with lightning deconvolution (UCSD School of Medicine Microscopy Core). Neuroblastoma cell lines were imaged with a Leica TCS SP5 microscope, HCC PL APO lambda blue 63x 1.4 oil lens.

### Neuroblastoma patient tissue FISH

FISH analysis was performed on 4 µm sections of FFPE blocks. Slides were deparaffinized, dehydrated and incubated in pre-treatment solution (Dako, Denmark) for 10 min at 95–99°C. Samples were treated with pepsin solution for 2 min at 37°C. For hybridization, the Zyto*Light* ^®^ SPEC MYCN/2q11 Dual Color Probe (ZytoVision, Bremerhaven, Germany) was used. Incubation took place overnight at 37°C, followed by counterstaining with 4,6-diamidino-2-phenylindole (DAPI). For each case, signals were counted in 50 non-overlapping tumour cells using a fluorescence microscope (BX63 Automated Fluorescence Microscope, Olympus Corporation, Tokyo, Japan). Computer-based documentation and image analysis was performed with the SoloWeb imaging system (BioView Ltd, Israel) MYCN amplification (MYCN FISH+) was defined as MYCN/2q11.2 ratio > 4.0, as described in the INRG report^27^.

### Quantification of FISH foci

Quantification of FISH foci was performed using the ImageJ-Find maxima function in a supervised fashion. For quantification of pixel intensity, the ImageJ-Pixel intensity function was used. These two GBM patient tissue FISH images were obtained as part of a phase II lapatinib GBM clinical trial described previously. In brief, patients were administered 750 mg of lapatinib orally twice a day (BID) for 7 to10 days (depending on whether treatment interval fell over a weekend) before surgery, the time to steady state. Blood and tissue samples were obtained at the time of resection^2^.

### Construction of PC3-TetO cell line

The insertion of tetO repeats was conducted through CRISPR/cas9 mediated approaches. And the plasmids: pSP2-96-mer TetO-EFS-BlaR and F9-TetR-EGFP-IRES-PuroR used in this section were kind gifts from Dr. Huimin Zhao^21^. Briefly, the intergenic region between MYC and PVT1 was selected as the insertion region on the basis that it is amplified in PC3 cells on ecDNA with high frequency. DNA sequences were retrieved from UCSC Genome Brower, repetitive and low complexity DNA sequences were annotated and masked by RepeatMasker in the UCSC Genome Browser. The guide sequences of sgRNAs were designed by CRISPRdirect web tool^28^, and their amplification was confirmed with WGS data. The guide sequence selected was constructed into pSpCas9(BB)-2A-Puro (PX459) [pSpCas9(BB)-2A-Puro(PX459) was a gift from Feng Zhang (Addgene plasmid #62988; http://n2t.net/addgene:62988; RRID: Addgene_62988)]. Repair donor was obtained through PCR amplification, using pSP2-96-merTetO-EFS-BlaR plasmid as template, as well as primers containing the 50nt homology arm upstream and downstream of the predicted cutting site.

The transfection of CRISPR/Cas9 plasmid and 96-mer TetO EGFP-BlastR donor into PC3 cells was conducted with X-tremeGENE HP transfection reagent according to manufactory instruction with CRISPR/Cas9 plasmid only or 96-mer TetO EGFP-BlastR only using as negative control. 2 days after transfection, Blasticidin was added to the culture medium for 3 days, at a time point that the majority of the cells in the negative control groups have died while more cells survived in the group with transfection of CRISPR/Cas9 plasmid and donor. The surviving cells were subjected to limited dilution in 96-well plate, with Blasticidin being added all the time. Surviving clones were expanded and their genomic DNA were extracted and subjected to genotyping with a pair of primer flanking the inserted region. PCR product of genotyping results were subjected to sanger sequencing to confirm the insertion at predicted cutting site. Clones with positive genotyping band will be expanded and metaphase cells were collected. Double FISH with FISH probe against Tet operator and against MYC FISH probe was performed on metaphase spread. PC3 cells with TetO repeats were infected with lentivirus containing the F9-TetR-EGFP-IRES-PuroR, and 2 days after infection puromycin was added into culture medium to establish a stable cell line that is able to image ecDNA with the aid of EGFP visualization.

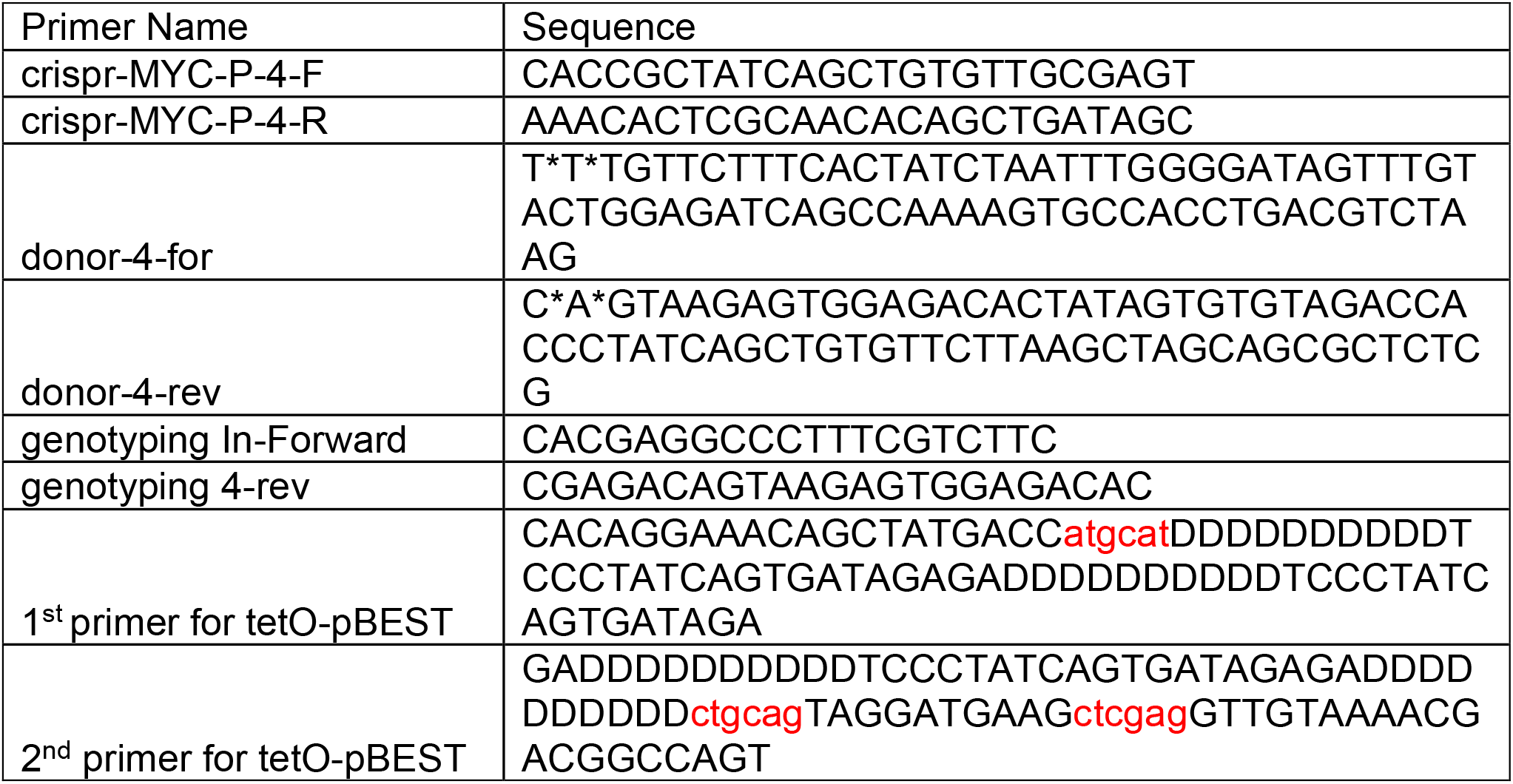

### Live cell imaging of ecDNA

PC3 TetO TetR-GFP cell line was transfected with PiggyBac vector expressing H2B-SNAPf and the super PiggyBac transposase (2:1 ratio) as previously described^29^. Stable transfectants were selected by 500µg/ml G418 and sorted by flow cytometry. To facilitate long-term time lapse imaging, 10µg/ml human fibronectin was coated in each well of 8-well lab-tek chambered cover glass. Prior to imaging, cells were stained with 25nM SNAP tag ligand JF_669_^22^ at 37°C for 30 minutes followed by 3 washed with regular medium for 30 minutes total. Cells were then transferred to an imaging buffer containing 20% serum in 1x Opti-Klear live cell imaging buffer at 37°C. Cells were imaged on a Zeiss LSM880 microscope pre-stabilized at 37°C for 2 hours. We illuminated the sample with 1.5% 488nm laser and 0.75% 633nm laser with the EC Plan-Neofluar 40x/1.30 oil lens, beam splitter MBS 488/561/633 and filters BP 495-550 + LP 570. Z-stacks were acquired with 0.3µm z step size with 4-minute intervals between each volumetric imaging for a total of 16 hours.

### Colony formation assay

TR-14 cells were taken from 60 days of treatment with either DMSO, 50 nM Palbociclib, or 5 nM Abemaciclib, and seeded into a poly-D-lysine coated 24-well plate at 20,000 cells per well. After 24 h, the cells from each condition were treated with either DMSO, 50 nM Palbociclib, or 5 nM Abemaciclib over 20 days, in triplicate. At 20 days, crystal violet staining procedure was performed. Briefly, cell culture media was aspirated, cells were washed gently with PBS, fixed in 4% paraformaldehyde in PBS for 20 min, stained with 2 mL of crystal violet solution (50 mg in 50 mL 10% ethanol in MilliQ water), washed 1x with PBS, and dried for 30 min. The area intensity was calculated using the ColonyArea plugin in ImageJ^30^.

### CellTiter-Glo

TR-14 cells were taken from 60 days of treatment with either DMSO, 50 nM Palbociclib, or 5 nM Abemaciclib and seeded into white flat-bottom 96 WPs (Corning) in 100 µl media at a density of 500 cells/well. After 24 h, the cells were treated with either vehicle, 50 nM Palbociclib, or 5 nM Abemaciclib (50 µL of drug solution/well). Cell viability was determined using CellTiter-Glo Luminescent Cell Viability Assay (Promega) at 3, 6, and 9 days after drug was added, following the manufacturer’s protocol.

### Immunoblotting

Whole-cell protein lysates were prepared by lysing cells in Silly lysis buffer. Protein concentrations were determined by bicinchoninic acid assay (BCA, Thermo Fisher). 10 µg of protein were denatured in Laemmli buffer at 95 °C for 5 minutes and 1mM DTT was added. Lysates were loaded onto 10% Tris-Glycin (Thermo Fisher) for gel electrophoresis. Proteins were transferred onto Immobilon-FL PVDF membranes (Sigma Aldrich), blocked Odyssey Blocking Buffer in TBS for 1 hour and incubated with primary antibodies overnight at 4°C, then secondary antibodies for 1 hour at room temperature. Fluorescent signal was detected using the Odyssey CLx imaging system. Quantification was performed with LI-COR Image Studio Software.

### Flow cytometry

Single cell suspensions were made and passed through a cell filter to ensure single cell suspension. Cells were suspended in flow cytometry buffer (HBSS buffer without calcium and magnesium, 1x Glutamax, 0.5% (v/v) FBS, 10mM HEPES). EGFRvIII mab 806^31^ was added at 1ug per million cells and incubated on ice for one hour. Cells were washed in flow cytometry buffer and resuspended in buffer with anti-mouse alexa-488 antibody (1:1000) for 45 minutes on ice in the dark. Cells were washed again with flow cytometry buffer and resuspended in flow cytometry buffer at approximately 4 million cells per milliliter. Cells were sorted using a Sony SH800 FACS sorter and was calibrated and gating was informed using a secondary only negative control.

### Quantitative PCR (qPCR)

DNA extraction was performed using the NucleoSpin Tissue kit (Macherey-Nagel), following the manufacturer’s protocol. qPCR was performed using 50 ng or 1.5 µl of template DNA and 0.5 µM primers with SYBR Green PCR Master Mix (Thermo Fisher Scientific) in FrameStar 96-well PCR plates (4titude). Reactions were run and monitored on a StepOnePlus Real-Time PCR System (Thermo Fisher Scientific) and Ct values were calculated with the StepOne Plus software v.2.3 (Thermo Fisher Scientific).

CDK4 Fwd: AAAGTTACCACCACACCCCC

CDK4 Rev: AGTGCTAAGAAAGCGGCACT

### Quantification of single cell ecDNA segregation patterns

We generate the theoretically expected distribution of ecDNA copy number fractions after a single cell division under different models of ecDNA segregation by stochastic computer simulations implemented in C++. Briefly a single cell is initiated with a random number of ecDNA copies *n*, drawn from a uniform distribution *U*(20,200). EcDNA is amplified and 2*n* ecDNA copies are segregated between two daughter cells following a Binomial trial *B*(2*n, p*), with segregation probability *p*. Here, *p* = 1/2 corresponds to random segregation and *p* > 1/2 to a biased random segregation. This results in two daughter cells with ecDNA copy number *n*_1_∼*B*(2*n, p*) and *n*_2_ = *n* − *n*_1_. The fraction of segregated ecDNA *f* is then calculated via

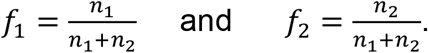

Iterating the process 10^7^times generates the expected distribution of *f* as shown in Figure 1c. Similarly, we can generate an expected distribution of *f* for chromosomal patterns of inheritance. For perfect chromosomal segregation, we have *f*_1_ = *f*_2_ = 1/2. To allow for mis-segregation we introduce a probability *u* = 0.05 such that *n*_1_ = *n* ± 1 and *n*_2_ = *n* − *n*_1_. We use Kolmogorov-Smirnov statistics to compare the theoretically expected and experimentally observed distributions of ecDNA copy number fractions under these different scenarios.

### Stochastic simulations of ecDNA population dynamics

We implemented individual based stochastic computer simulations of the ecDNA population dynamics in C++. For each cell, the exact number of ecDNA copies is recorded through the simulation. Cells are chosen randomly but proportional to fitness for proliferation using a Gillespie algorithm. The simulation is initiated with one cell carrying *n*_0_ copies of ecDNA. The proliferation rate of cells without ecDNA is set to *r*^−^ = 1 (time is measured in generations). A fitness effect for cells with ecDNA then corresponds to a proliferation rate *r*^+^ = *s*. Here, *s* > 1 models a fitness advantage, 0 < *s* < 1 a fitness disadvantage and *s* = 1 corresponds to no fitness difference (neutral dynamics, *r*^+^ = *r*^−^).

During proliferation, the number of ecDNA copies in that cell are doubled and randomly distributed into both daughter cells according to a Binomial trail *B*(*n, p*) with success rate *p* = 1/2. If a cell carries no ecDNA, no daughter cell inherits ecDNA. We terminate simulations at a specified cell population size. We output the copy number of ecDNA for each cell at the end of each simulation, which allows us to construct other quantities of interest, such as the ecDNA copy number distribution, the time dynamics of moments, the power law scaling of tails or the Shannon diversity index. We use Kolmogorov-Smirnov statistics to test similarity between simulated and experimental ecDNA copy number distributions and Shapiro-Wilk statistics to test for deviations from normality.

### Sampling and resolution limits

We ran an *in-silico* trial to test our ability to reconstruct the true ecDNA copy number distribution from a sampled subset of varying sizes. We constructed a simulated ecDNA copy number distribution from 2 × 10^6^ cells using our stochastic simulations. We then performed 500 random samples of 25, 50, 100 and 500 cells, reconstructed the sampled ecDNA copy number distribution and compared similarity to the true copy number distribution using Kolmogorov-Smirnov statistics. The distribution converges to the true distribution with increasing sampling size and a comparably small sample of 100 to 500 cells is sufficient to reconstruct the true underlying ecDNA copy number distribution.

### Mathematical description of ecDNA

#### dynamics Deterministic two population model without selection

In the simplest representation of the model, we discriminate cells that do or do not carry copies of ecDNA. We denote cells with copies of ecDNA as *N*^+^(*t*) and cells without copies of ecDNA with *N*^−^(*t*). We can write for the change of these cells in time *t*

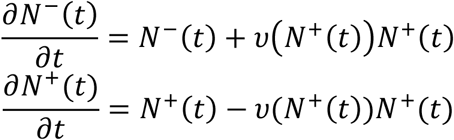

where *v*(*N*^+^(*t*)) corresponds to the loss rate of random complete asymmetric ecDNA segregation. We find for the fraction of cells carrying ecDNA *f*^+^(*t*) in an exponentially growing population

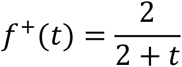

The fraction of cells carrying ecDNA decreases with ∼1/*t* if ecDNA is neutral. Thus, copies of neutral ecDNA are only present in a small subpopulation of tumour cells.

#### Deterministic two population model with selection

Above equations can be modified to allow for a fitness advantage *s* > 1 for cells carrying ecDNA.

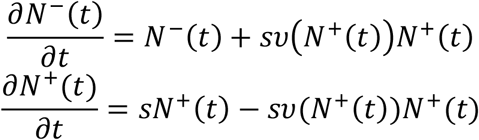

The solution to this set of equations is

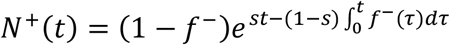

In the case of positive selection, the fraction of cells with ecDNA *f*^+^ → 1. For sufficiently long times, the tumour will be dominated by cells carrying ecDNA.

#### Stochastic dynamics of neutral ecDNA

We are also interested in the stochastic properties of ecDNA dynamics in a growing population. We therefore move to a more fine-grained picture and consider the number of cells *N*_*k*_ (*t*) with *k* copies of ecDNA at time *t*. The dynamic equation for neutral copies of ecDNA becomes

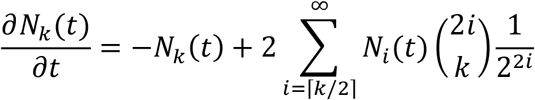

It is more convenient to work with the cell density *ρ* instead of cell numbers *N*. Normalizing above equation, we get for the density *ρ*_*k*_ of cells with *k* ecDNA copies

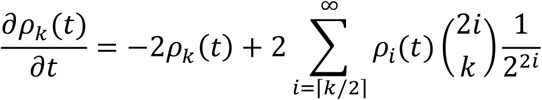

#### Moment dynamics for neutral ecDNA copies

With above equation for the density of cells with *k* ecDNA copies, we can calculate the moments of the underlying probability density function. In general, the *l*-th moment can by calculated via

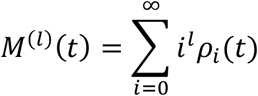

It can be shown that all moments scale with *M*^(*l*)^(*t*)∼*t*^*l*−1^ and we find explicitly for the first two moments

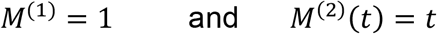

The mean ecDNA copy number in an exponentially growing population remains constant for neutral ecDNA copies. The variance of the ecDNA copy number increases linearly in time.

#### Stochastic dynamics of ecDNA under positive selection

Above equations can be generalized to accommodate positive selection (*s* > 1) for ecDNA copies. The set of dynamical equations for cell densities becomes

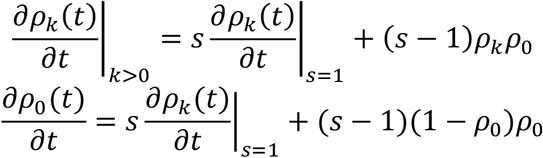

A general solution to these equations is challenging, but still important quantities, e.g., the moment dynamics and the scaling behavior can be calculated explicitly.

#### Moment dynamics for ecDNA under positive selection

A generalized equation for the dynamics of moments directly follows from above equations. We have

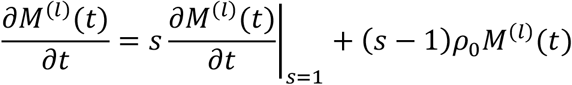

This implies for the first moment 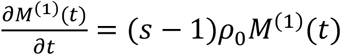, which then can be solved for the first moment

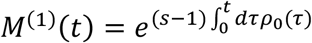

Similarly, the dynamic equation for the second moment becomes 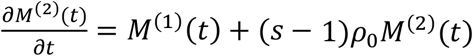 and we find

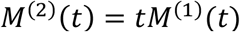

The first moment increases exponentially initially. However, with increasing mean copy number, the rate of cells transition into a state without ecDNA is decreasing and the increase of the mean ecDNA copy number slowly levels of. Note, for *s* = 1 we recover the previous results for the moments of neutral ecDNA amplifications.

#### Scaling wave solution and limiting behavior of the ecDNA copy number distribution

In the following, we are interested in the scaling behavior of the ecDNA copy number distribution. Our general time dynamics considers discrete copy number states. To make further analytical progress, we now consider continues states in the following calculations. This is an approximation that works well for the case of many ecDNA copies, but might be inaccurate for cells with very few copies of ecDNA. Under this continues assumption, the change of the ecDNA copy number distribution becomes

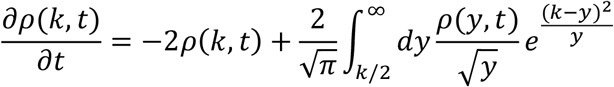

Here, we also replaced the Binomial by a Normal distribution. Given the exponential character of the ecDNA distribution, we proceed with an Ansatz in the form of a scaling wave solution

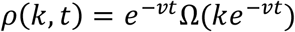

Plugging this Ansatz into the dynamical equation and taking all terms into careful consideration, it follows that 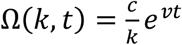, where *c* is an undetermined constant and thus we have

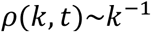

For sufficiently large ecDNA copy number. With other words, we expect the right-hand tail of the ecDNA copy number distribution to follow a power law proportional to the inverse of the ecDNA copy number. This prediction is confirmed in stochastic computer simulations and can also be observed in experimentally measured distributions.

#### Genome editing using CRISPR-Cas9 ribonucleoprotein

Genome editing in COLO320-DM cells were performed using Alt-R S.p. Cas9 Nuclease V3 (IDT, Cat# 1081058) complexed with sgRNA (Synthego) following Synthego’s RNP transfection protocol using the Neon Transfection System (ThermoFisher, Cat# MPK5000). Briefly, 10 pmol Cas9 protein and 60 pmol sgRNA for each 10 ul reaction were incubated in Neon Buffer R for 10 minutes at room temperature. Cell were washed with 1X PBS, resuspended in Buffer R, and 200,000 cells were mixed with for the pre-incubated ribonucleoprotein complex for each 10 ul reaction. The cell mixture was electroporated following the manufacturer’s protocol using the following settings: 1700 V, 20 ms, 1 pulse. Cells were cultured for 10 days afterwards; cell counts and ecDNA copy number data were collected at day 3, 6, and 10. To estimated ecDNA copy numbers, we performed metaphase chromosome spreading followed by FISH as described above. All sgRNA sequences are in table below.

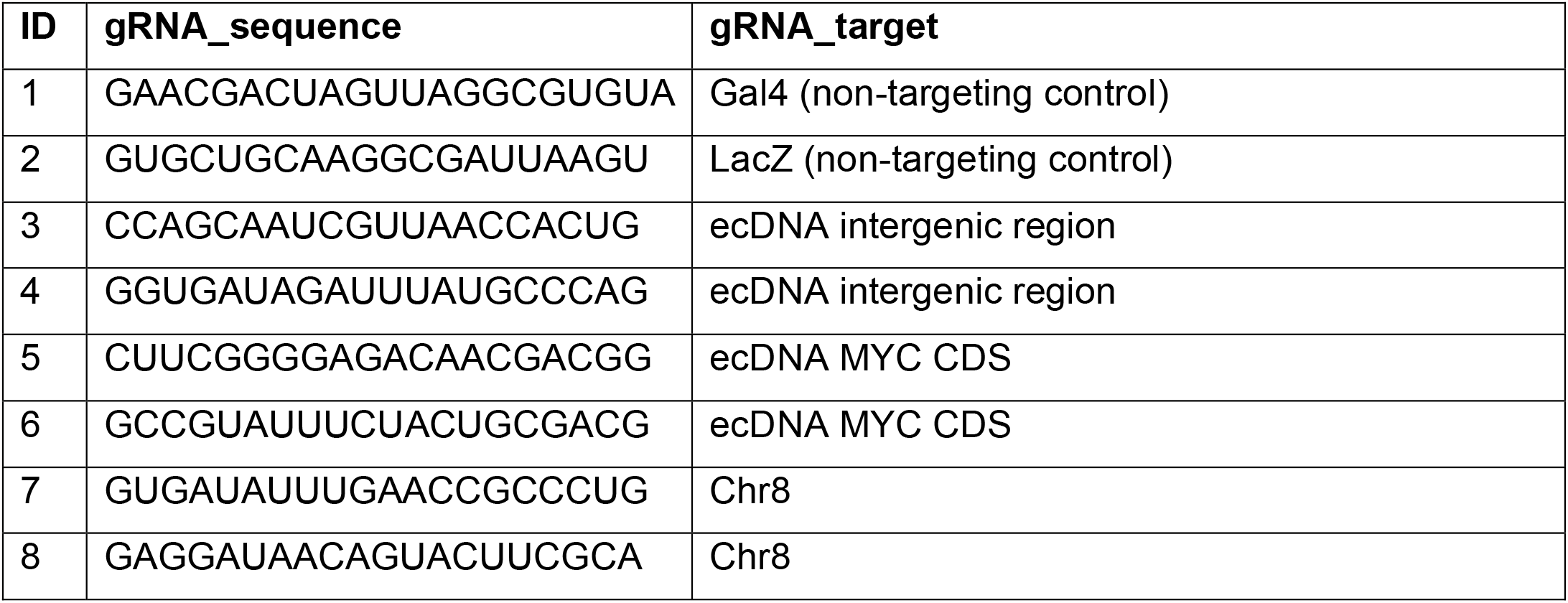

#### FISH probes

ZytoLight SPEC CDK4/CEN 12 Dual Color Probe (ZytoVision)

Zyto*Light* SPEC MYCN/2q11 Dual Color Probe (ZytoVision)

Empire Genomics EGFR FISH Probe

Empire Genomics MYC FISH Probe

Empire Genomics FGFR2 FISH Probe

#### Antibodies

β-Actin (8H10D10) Mouse mAb #3700 (Cell Signaling)

CDK4 (D9G3E) Rabbit mAb #12790 (Cell Signaling)

IRDye 780RD Secondary Antibody (Licor)

IRDye 800CW Secondary Antibody (Licor)

Aurora B Polyclonal Antibody #A300-431A (ThermoFisher)

