## Supplementary Information for "Principles of ecDNA random inheritance drive rapid genome change and therapy resistance in human cancers"

Mathematical modelling of ecDNA dynamics

### Introduction

Here we develop and analyse a baseline mathematical model of random ecDNA segregation in exponentially growing tumour populations. This will allow us to work out a set of dynamic predictions to distinguish ecDNA behaviour under neutral or positive selection. We will see that certain properties, such as the mean ecDNA copy number per cell and the fraction of cells with and without ecDNA fundamentally differ between these two scenarios.

We first present stochastic computer simulations using an agent-based model and compare the simulations against experimental data. Next, we develop a complete and fine-grained picture of ecDNA dynamics and work out the theoretical dynamics of moments as well as the expected scaling of the ecDNA copy number distribution. This is followed by a simplified deterministic approximation that will allow us to follow the change of cell populations with and without ecDNA copies in time. These analytical results are compared both against experimental data as well as stochastic simulations.

Our mathematical model is based on five assumptions: (i) ecDNA copies are segregated randomly between daughter cells; (ii) the cell population is exponentially growing; (iii) ecDNA replicates at the same rate as chromosomal DNA doubling during the cell cycle; (iv) the population starts with a single cell carrying a single copy of ecDNA; (v) a cell that has lost all ecDNA does not regain them.

Our reasoning for these assumptions is as follows: (i) We have experimentally verified this property across different cell lines with different ecDNA amplified genes. This is the fundamental dynamic property that distinguishes ecDNA copy number evolution from the evolution of somatic copy number alterations on chromosomes. (ii) We are interested in ecDNA evolution in growing tumour populations. (iii) This assumption can be justified retrospectively. If ecDNA is amplified with any coefficient $> 2$ the ecDNA copy number per cell explodes within a few generations and each cell would be expected to carry thousands of ecDNA copies. This ecDNA copy number inflation is not observed in any of the cell line or patient data. (iv) Here we are interested in specific types of ecDNA amplifications. If we say a cell carries $k$ copies of ecDNA, we mean exactly $k$ copies of one particular complex amplification, e.g. EGFR in Glioblastoma or MYCN in Neuroblastoma. These are large and complex genomic structures, and we assume that their origin is a single catastrophic event in the evolutionary history of a tumour and a repeated production of the exact same circular DNA structure containing millions of base pairs is extremely unlikely. There very well can be situations, where cells carry multiple types (species) of ecDNA, e.g. an EGFR and MYC amplification. In this situation, we would introduce two copy numbers $k_1$ and $k_2$ that keep track of the temporal evolution of those two species independently. (v) ecDNA formation is a rare, random event. Most ecDNA impose a metabolic load on the cell, are deleterious to its fitness and lost rapidly. Very rarely, an ecDNA is created that carries a proliferative element (e.g. an oncogene) which provides a growth and proliferative advantage to the cell.

Our notation will be as follows. $N(t)$ refers to the number of cells $N$ at any particular time $t$ during the growth of the tumour. $N_k(t)$ refers to the number of cells with exactly $k$ copies of ecDNA at time $t$. The copy number per cell, $k$, can in principle range from zero to infinity. With this we can formulate the equation for the expected temporal change of cells with $k$ ecDNA copies. For simplicity, we first explain the case of neutral ecDNA dynamics (cell with and without ecDNA have the same properties).

### 1.1 Agent based stochastic computer simulations of ecDNA segregation

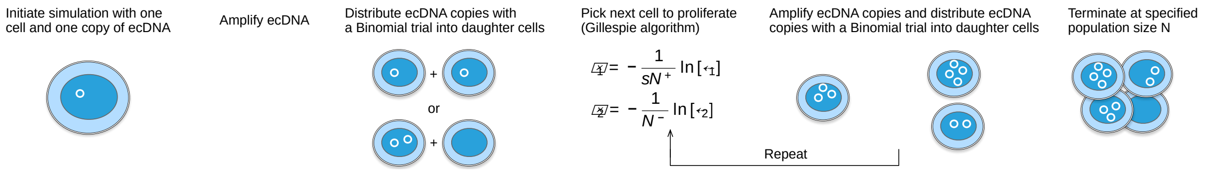


**Figure SI 1.** Schematic of the stochastic simulations for random ecDNA segregation in exponentially growing tumour populations.

A schematic of the simulations can be found in Figure SI 1. All simulations are exact agent-based implementations of the underlying stochastic process. Simulations are initiated with a single cell carrying a single copy of ecDNA. Upon proliferation, the number of ecDNA copies in a cell are doubled and distributed between two daughter cells following a Binomial trial with success probability $1/2$. From thereon, the next cell to proliferate is chosen following a Gillespie algorithm. Briefly, we draw two random numbers $\zeta_1$ and $\zeta_2$ from a Uniform distribution in the interval $[0,1]$ and calculate the corresponding reaction times for cells with ecDNA ($N^+$) and cells without ecDNA ($N^-$), given by $\tau_1 = -\frac{1}{sN^+}\ln[\zeta_1]$ and $\tau_2 = -\frac{1}{N^-}\ln[\zeta_2]$. Whichever reaction time is smaller, is the next cell chosen for proliferation. Again, the ecDNA copy number of the cell is doubled and distributed into two daughter cells following a Binomial trial with success rate $1/2$. This process is iterated until the cell population reaches a predefined number of cells $N$. The same stochastic process can be used to simulate related dynamics for non-random ecDNA segregation. We just need to replace the Binomial trial by a segregation probability of interest, e.g. we could have non-random biased segregation with $p > 1/2$, or strict chromosomal segregation where each daughter cell always receives equal number of ecDNA copies. These simulations introduce two sources of stochasticity. The next cell to proliferate is picked at random, but proportional to fitness. The Gillespie algorithm (Gillespie, Journal of Physical Chemistry 1977) offers an exact stochastic implementation of the underlying Markov Chain and its implementation is standard in these types of individual based simulations. The second source of randomness emerges from the segregation of ecDNA copies into daughter cells after division. Computer simulations of (non)random ecDNA segregation have been implemented in C++ and the code to run the simulations is available https://github.com/BenWernerScripts.

### 1.2 Comparison of stochastic simulations and experimental observations

The final output of our stochastic simulations is a population of cells, each cell with a particular ecDNA copy number. These copy number distributions can be followed over time, and all information of interest, e.g. the population of cells with and without ecDNA, the mean and variance of the ecDNA distribution, the actual ecDNA copy number distribution as well as the power law scaling of the ecDNA distribution can be constructed.

We use a Kolmogorov-Smirnov test to compare the ecDNA copy number distributions from stochastic computer simulations and experimentally observed distributions in patients or cell line experiments. The test first gives the $KS_d$ distance, with smaller values indicating better agreement. It also allows us to calculate a $p_{KS}$ value. The test compares two probability distributions for distance d, the $p$-value corresponds to the probability of obtaining d or smaller given the that the two distributions are different. For the ecDNA copy number distributions, we also use the Shapiro-Wilk statistics, to test for deviations from a normal distribution. In addition, to show goodness of fits, we added Quantile-Quantile plots for all comparisons of experimental and theoretical distributions.

| Sample | $\mathbf{KS_d^{random}}$ | $p_{KS}^{\mathbf{random}}$ | $\mathbf{KS_d^{non-random}}$ | $p_{KS}^{\mathbf{non-random}}$ | $p_{\mathbf{ShapiroWilk}}$ | #samples |
|---|---|---|---|---|---|---|
| PC3_Myc | 0.065 | 0.375 | 0.46 | 0 | 0.758 | 200 |
| SNU16_Myc | 0.039 | 0.918 | 0.49 | 0 | 0.939 | 194 |
| SNU16_fgfr2 | 0.063 | 0.415 | 0.49 | 0 | $4.2 \times 10^{-9}$ | 196 |
| GBM39_EGFR | 0.072 | 0.221 | 0.46 | 0 | 0.001 | 210 |
| COLO_Myc | 0.033 | 0.973 | 1 | 0 | 0.196 | 206 |

**Table SI 1.** Test statistics to compare the theoretical distributions with experimental observations for the single cell ecDNA segregation probabilities as presented in Figure 1c in the main text. The similarity of the two distributions is tested by a Kolmogorov-Smirnov test for two competing hypothesis, random ecDNA segregation and non-random chromosomal segregation. We also test for normality using the Shapiro-Wilk statistics.

| Sample | $\mathbf{KS_d^{random}}$ | $p_{KS}^{\mathbf{random}}$ | $\mathbf{KS_d^{non-random}}$ | $p_{KS}^{\mathbf{non-random}}$ | $p_{\mathbf{ShapiroWilk}}$ | #samples |
|---|---|---|---|---|---|---|
| PC3_Myc | 0.091 | 0.074 | 0.986 | 0 | $3.1 \times 10^{-13}$ | 200 |
| SNU16_Myc | 0.052 | 0.662 | 0.999 | 0 | $9.9 \times 10^{-4}$ | 194 |
| SNU16_fgfr2 | 0.066 | 0.359 | 1 | 0 | $1.9 \times 10^{-12}$ | 196 |
| GBM39_EGFR | 0.071 | 0.237 | 0.977 | 0 | $6.6 \times 10^{-9}$ | 210 |
| COLO_Myc | 0.075 | 0.196 | 0.994 | 0 | $2.4 \times 10^{-11}$ | 206 |
| GBM1 | 0.141 | 0.073 | 0.882 | 0 | 0.019 | 85 |
| GBM2 | 0.082 | 0.914 | 0.757 | 0 | 0.028 | 46 |
| GBM3 | 0.138 | 0.131 | 0.843 | 0 | 0.003 | 72 |
| GBM4 | 0.254 | 0.004 | 0.759 | 0 | 0.014 | 101 |
| GBM5 | 0.163 | 0.01 | 0.831 | 0 | 0.004 | 103 |
| GBM6 | 0.159 | 0.124 | 0.833 | 0 | 0.057 | 55 |
| Chp212 | 0.193 | 0.048 | 0.963 | 0 | $1.2 \times 10^{-10}$ | 154 |
| TR14_MYCN | 0.047 | 0.681 | 0.987 | 0 | $1.6 \times 10^{-8}$ | 232 |
| TR14_CDK4 | 0.091 | 0.174 | 0.855 | 0 | $1.7 \times 10^{-13}$ | 284 |
| NB4 | 0.098 | 0.177 | 1 | 0 | $1.2 \times 10^{-8}$ | 126 |
| NB7 | 0.129 | 0.313 | 0.999 | 0 | $4.8 \times 10^{-3}$ | 56 |
| NB8 | 0.074 | 0.375 | 0.983 | 0 | $1.3 \times 10^{-6}$ | 151 |
| NB10 | 0.176 | 0.004 | 0.996 | 0 | $3.3 \times 10^{-6}$ | 98 |
| NB13 | 0.271 | 0.001 | 0.999 | 0 | 0.004 | 155 |

**Table SI 2.** Test statistics to compare the theoretical ecDNA copy number distributions with experimental measured ecDNA copy number distributions in patient and cell line data as presented in Figure 2 b and c in the main text. The similarity of the two distributions is tested by a Kolmogorov-Smirnov test for two competing hypothesis, random ecDNA segregation and non-random chromosomal segregation. We also test for normality using the Shapiro-Wilk statistics.

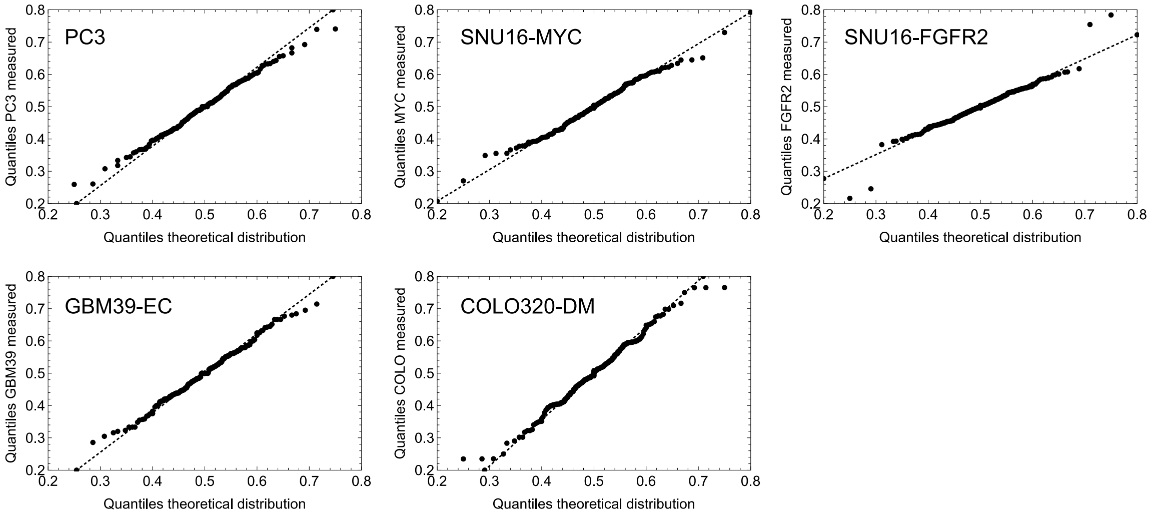


**Figure SI 2.** Quantile-Quantile_plots to compare the theoretical and experimental distributions from single cell ecDNA segregation probabilities presented in Figure 1c in the main text.

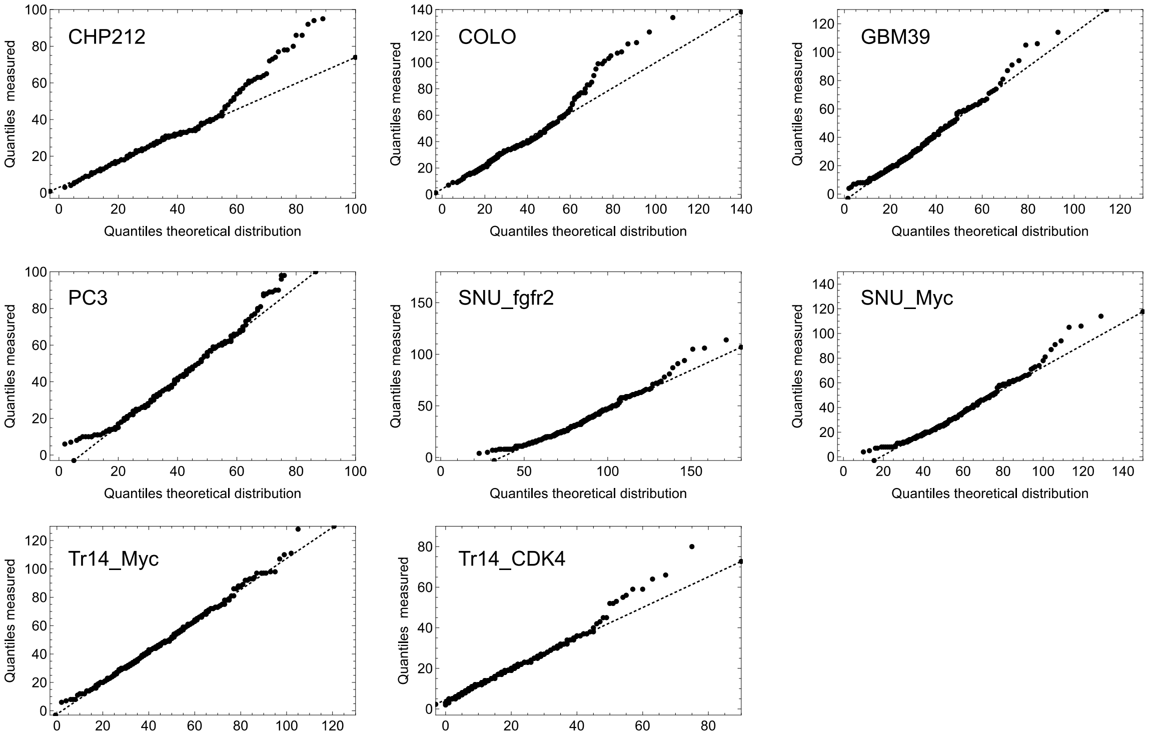


**Figure SI 3.** Quantile-Quantile_plots to compare the theoretical and experimental ecDNA copy number distribution in cell lines presented in Figure 2b in the main text.

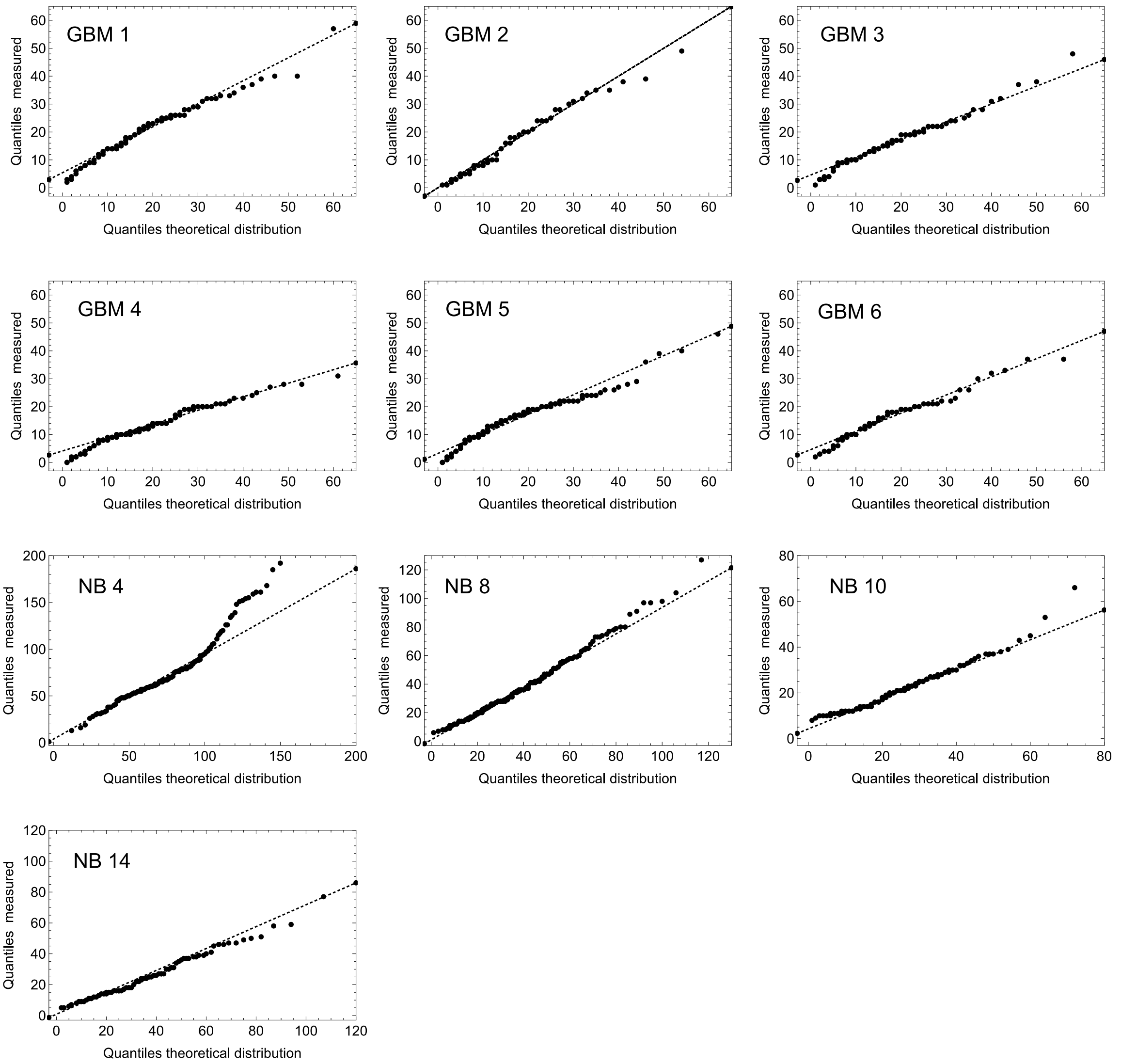


**Figure SI 4.** Quantile-Quantile_plots to compare the theoretical and experimental ecDNA copy number distribution in samples of Glioblastoma and Neuroblastoma patients presented in Figure 2c in the main text.

### 1.3 Finite sampling and resolution limits

In our stochastic simulations, we have the freedom to in principal sample and analyse as many single cell ecDNA copy number profiles as we want. This is obviously not the case in our experimental data due to technical and financial limitations. We thus tested if we can reconstruct the ecDNA copy number distribution with limited single cell resolutions. We generated a distribution of ecDNA copy numbers by simulating a tumour with $10^7$ cells and ecDNA under positive selection $s = 2$. We then sampled 10,000 times 25, 50, 100 and 500 cells respectively, constructed the ecDNA copy number distribution and calculated the Kolmogorov distance

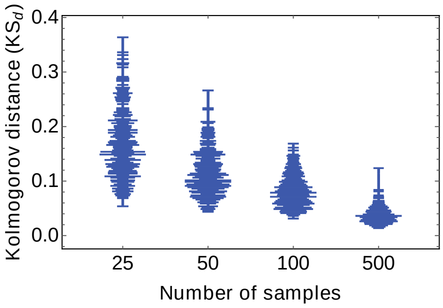


**Figure SI 5.** Sampling of the ecDNA copy number distribution. We took $10^4$ samples with $25, 50, 100$ and $500$ cells respectively from a single simulation of the ecDNA distribution of $10^7$ cells. Shown are the corresponding distributions of Kolmogorov distances. Resolution increases with sample size. Kolmogorov distances for samples of 100 cells are comparable to our experimental observations.

of the sampled distribution to the true (non-sampled) distribution. As expected, the resolution increases with sample size. More importantly, we find Kolmogorov distances that are comparable to experimental data comparisons and a sample size in the order of 100 cells already allows us to capture important aspects of the ecDNA copy number distribution.

### 2.1 Stochastic dynamics of ecDNA copy numbers under neutral selection

The dynamic equation for the number of cells $N_k(t)$ with $k$ neutral copies of ecDNA with time $t$ becomes

$$\frac{\mathrm{d}N_k(t)}{\mathrm{d}t} = -N_k(t) + 2\sum_{i=\lceil k/2\rceil}^{\infty} N_i(t)\binom{2i}{k}\frac{1}{2^{2i}}$$

This is a set of, in principal, infinitely many coupled differential equations, formally known as the Master equation of the underlying Markovian stochastic process. It describes the evolution of all states the system at question can be in. In our case, all possible states correspond to the number of cells with $k$ copies of ecDNA. The left-hand side is the time derivative of the number of cells with $k$ ecDNA copies. The right-hand side collects all possible events (rates) that change this number. If a cell with $k$ copies divides, its copies are amplified and randomly distributed between both daughter cells. This reduces the number of cells with exactly $k$ copies, reflected by the first term $-N_k(t)$. The second term on the right-hand side of the equation collects all cells of the system that gain $k$ copies of ecDNA due to random segregation amongst daughter cells. Upon cell proliferation, $2i$ copies are randomly segregated amongst two daughter cells. The number of ecDNA copies $k$ in a daughter cell follows a Binomial distribution with success rate $1/2$

$$B\left(k \,\middle|\, n = 2i, p = \frac{1}{2}\right) = \binom{2i}{k}\frac{1}{2^{2i}}$$

It turns out that working with cell densities $\rho_k$rather than total cell numbers $N_k$ is advantageous. We therefore decouple population growth and demographic changes and write $N_k(t) = N(t)\rho_k(t)$ with $\sum_{i=1}^{\infty}\rho_i(t) = 1$ and $N(t) = \sum_k N_k(t)$ denotes the total number of cells at time t. We first can check that the structure of our equations is correct and we recover an exponentially growing population for $N(t)$ as we have claimed in our initial assumptions. We can write:

$$\frac{\mathrm{d}N(t)}{\mathrm{d}t} = -N(t) + 2N(t) \sum_{k=0}^{\infty} \sum_{i=\left\lceil \frac{k}{2} \right\rceil}^{\infty} \rho_i(t) \binom{2i}{k} \frac{1}{2^{2i}}$$

$$= -N(t) + 2N(t) \sum_{i=0}^{\infty} \rho_i(t) \frac{1}{2^{2i}} \sum_{k=0}^{2i} \binom{2i}{k}$$

$$= -N(t) + 2N(t) \sum_{i=0}^{\infty} \rho_i(t) \frac{1}{2^{2i}} 2^{2i} = N(t)$$

And we do find that the total population grows exponentially in time $N(t) = N(0)e^t$. This allows us to write for the temporal change of cell densities $\rho_k$ with $k$ ecDNA copy numbers:

$$\frac{\mathrm{d}\rho_k(t)}{\mathrm{d}t} = -2\rho_k(t) + 2 \sum_{i=\lceil k/2 \rceil}^{\infty} \rho_i(t) \binom{2i}{k} \frac{1}{2^{2i}}$$

### <u>2.2 Dynamics of Moments of ecDNA copies under neutral selection</u>

The Master equations above describe the full dynamics of the probability densities of the ecDNA copy number distribution. They therefore encode in principle all properties of the underlying stochastic process. However, a complete analytical treatment is challenging. Nevertheless, many aspects of the system are analytically tractable. We first discuss the dynamics of the moments of the ecDNA copy number distribution. In particular we are interested in the first and second moment, as they are directly related to the mean ecDNA copy number per cell and the expected variance of the ecDNA copy number distribution.

With above equation for the density of cells with $k$ ecDNA copies, we can calculate the moments of the underlying probability density function. In general, the $l$-th moment is calculated via

$$M^{(l)}(t) = \sum_{i=0}^{\infty} i^l \rho_i(t)$$

The moment $M^{(0)}(t)$ is just the sum over the density and by definition constant. The first moment corresponds to the average number of ecDNA copies per cell and we can write:

$$\frac{\mathrm{d}M^{(1)}(t)}{\mathrm{d}t} = -2M^{(1)}(t) + \sum_{k=0}^{\infty} \sum_{i=\lceil k/2 \rceil}^{\infty} k\rho_i(t) \binom{2i}{k} \frac{1}{2^{2i}}$$

$$= -2M^{(1)}(t) + \sum_{i=0}^{\infty} \rho_i(t) \frac{1}{2^{2i}} \sum_{k=0}^{2i} k \binom{2i}{k}$$

$$= -2M^{(1)}(t) + \sum_{i=0}^{\infty} \rho_i(t) \frac{1}{2^{2i}} (2i) 2^{2i-1} = 0$$

We therefore find $M^{(1)}(t) = \text{const}$ and the constant is given by the initial conditions. In most cases discussed here, we will have $M^{(1)}(t) = M^{(1)}(t = 0) = 1$. In the case of neutral ecDNA dynamics starting from a single cell containing a single copy of ecDNA, on average the population maintains one copy of ecDNA per cell.

Next, we are interested in the second moment $M^{(2)}(t)$. Following our calculations for the first moment we can similarly write:

$$\frac{\mathrm{d}M^{(2)}(t)}{\mathrm{d}t} = -2M^{(2)}(t) + \sum_{k=0}^{\infty} \sum_{i=\lceil k/2 \rceil}^{\infty} k^2 \rho_i(t) \binom{2i}{k} \frac{1}{2^{2i}}$$

$$= -2M^{(2)}(t) + \sum_{i=0}^{\infty} \rho_i(t) \frac{1}{2^{2i}} \sum_{k=0}^{2i} k^2 \binom{2i}{k}$$

$$= -2M^{(2)}(t) + \sum_{i=0}^{\infty} \rho_i(t) \frac{1}{2^{2i}} (2i + (2i)^2) 2^{2i-2} = M^{(1)}(t)$$

With the initial conditions for the mean ecDNA copy numbers above we find the expression $M^{(2)}(t) = t + \text{const}$. The constant can be fixed by the realisation that the variance of the ecDNA copy number distribution at time $t = 0$ should equal $0$ and we get $\text{Var}(t = 0) = \text{const} - 1^2 = 0$, and therefore $\text{const} = 1$ and simply have that the variance increases linearly in time for neutral ecDNA copies, $\text{Var}(t) = t$.

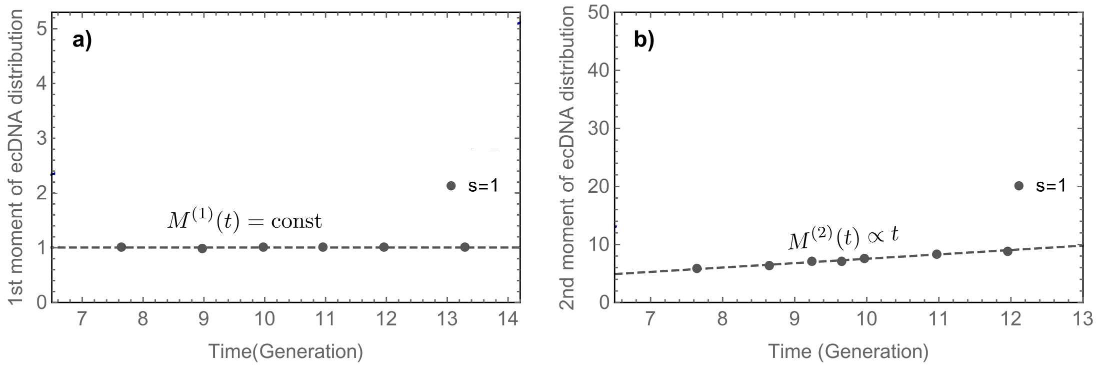


**Figure SI 6.** **a)** First and **b)** second moment of the ecDNA copy number distribution under neutral selection ($s = 1$). The mean number of ecDNA copies remains constant and the variance increases linearly in time. Stochastic simulations (points) are in very good agreement to theoretical predictions of polynomial increasing moments with time (dashed lines).

### 2.3 The scaling of the ecDNA copy number distribution in the continuous limit wave approximation

In the following, we are interested in the scaling behaviour of the ecDNA copy number distribution, e.g. what is the probability for a single cell to carry many copies of ecDNA. We will find that the right-hand tail of the ecDNA distribution towards large copy number scales with a power law inversely proportional to the copy number $k$.

Our general time dynamics describe discrete copy number states. To make further analytical progress, we now consider continuous states in the following calculations. This is an approximation that works well for the case of many ecDNA copies, but might be inaccurate for cells with very few copies of ecDNA. Under this continuous assumption, the change of the ecDNA copy number distribution becomes

$$\frac{\mathrm{d}\rho_k(t)}{\mathrm{d}t} = -2\rho(k,t) + \frac{2}{\sqrt{\pi}}\int_{k/2}^{\infty} \mathrm{d}y \frac{\rho_y(t)}{\sqrt{y}} e^{\frac{(k-y)^2}{y}}$$

Where we replaced the Binomial with a Normal distribution. Given the exponential character of the ecDNA distribution, we proceed with an Ansatz in the form of a scaling wave solution

$$\rho_k(t) = e^{-vt}\Omega(ke^{-vt}).$$

Plugging our ansatz into the differential equation for the density $\rho$ and setting $\frac{k}{2} \to 0$, we get

$$\frac{\mathrm{d}}{\mathrm{d}t}[e^{-vt}\Omega(ke^{-vt})] = -2e^{-vt}\Omega(ke^{-vt}) + \frac{2}{\sqrt{\pi}}\int_{k/2}^{\infty} \mathrm{d}y \frac{e^{-vt}\Omega(ke^{-vt})}{\sqrt{y}} e^{\frac{(k-y)^2}{y}}$$

With $z = ke^{-vt}$ and $v = 2$, this transforms into

$$-z\frac{\mathrm{d}}{\mathrm{d}z}\,\Omega(\mathrm{z}) = \frac{1}{\sqrt{\pi}}\int_{-\infty}^{\infty} \mathrm{d}m\,\Omega(\mathrm{z})\frac{1}{\sqrt{z}} e^{\frac{-m^2}{z}} = \Omega(\mathrm{z}).$$

This has the solution $\Omega(z) = c/z$ with an undetermined integration constant $c$. Plugging this back into our original ansatz and reversing all substitutions, this gives us for the scaling of the ecDNA copy number distribution

$$\rho_k(t) = \frac{c}{k}.$$

This predicts a power law scaling of the right-hand side tail of the ecDNA copy number distribution. Cells with very large copy number status become increasingly less likely for increasing $k$, for a sufficiently large tumour population, a considerable fraction of cells is expected to have large ecDNA copy number. This is indeed supported by observations both in cell line and patient data, where we recover these power law dependencies.

#### 3.1 Stochastic dynamics of ecDNA copies under constant positive selection

In the previous sections, we discussed the stochastic dynamics of extra-chromosomal DNA under neutral selection. In that scenario, ecDNA is present in cells, but does not change the proliferative fitness of the cell. Next, we consider the case of ecDNA that is under positive selection, or in other words, ecDNA that gives a positive fitness advantage to cells. This will be of particular interest to the dynamics and diversification of ecDNA in cancerous tissues.

In order to model a selection advantage, we introduce a selection coefficient $s > 0$. In this notation, $s = 1$ corresponds to neutral dynamics, $s > 1$ to a selection advantage of cells with ecDNA and $0 \leq s < 1$ to a selection disadvantage of cells without ecDNA. The Master equation then needs to be modified in the following way

$$\frac{\mathrm{d}N_k(t)}{\mathrm{d}t} = -sN_k(t) + 2s \sum_{i=\lceil k/2 \rceil}^{\infty} N_i(t) \binom{2i}{k} \frac{1}{2^{2i}}$$

$$\frac{\mathrm{d}N_0(t)}{\mathrm{d}t} = N_0(t) + 2s \sum_{i=1}^{\infty} N_i(t) \frac{1}{2^{2i}}$$

It can easily be checked that for $s \to 1$, we recover the Master equation in the neutral selection case. Above general Master equation for the selection case can also be written in a more compact form. Changing to the densities again, this compact form is given by

$$\left.\frac{\mathrm{d}\rho_k(t)}{\mathrm{d}t}\right|_{k>0} = s \left.\frac{\mathrm{d}\rho_k(t)}{\mathrm{d}t}\right|_{s=1} + (s-1)\rho_k\rho_0$$

$$\frac{\mathrm{d}\rho_0(t)}{\mathrm{d}t} = s \left.\frac{\mathrm{d}\rho_k(t)}{\mathrm{d}t}\right|_{s=1} + (s-1)(1-\rho_0)\rho_0$$

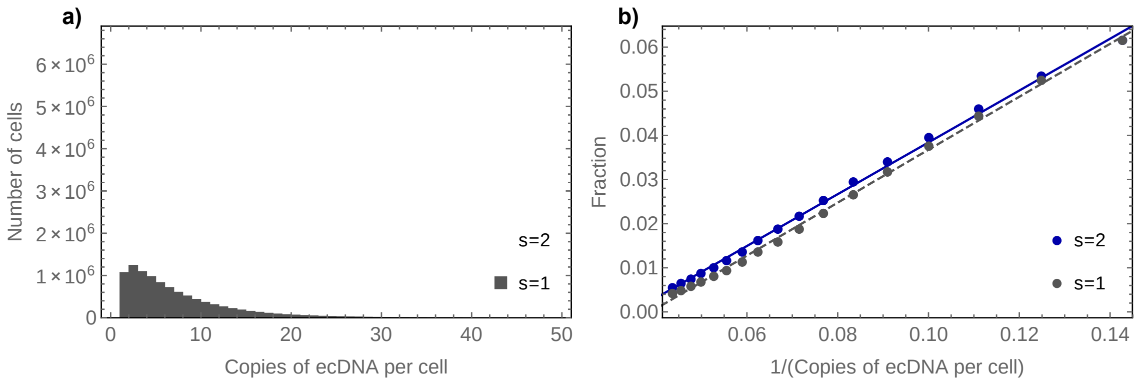


**Figure SI 7.** Distribution and scaling of the ecDNA copy number distribution. **a)** Distribution of the ecDNA copy number distribution for neutral (grey) and positively selected (blue) ecDNA evolution for 1000 repeats of stochastic simulations for tumours of $10^4$ cells. Overall, more cells carry copies of ecDNA if positively selected compared to the neutral case. **b)** The scaling of the right-hand tail of the ecDNA distribution follows the predicted 1/k scaling (dots = stochastic simulations, lines = theoretical expectation).

Allowing for selection adds an additional non-linear term to the original Master equation. We can also check the growth of the tumour population with ecDNA under positive selection. The equation for the total population now becomes

$$\frac{\mathrm{d}N(t)}{\mathrm{d}t} = sN(t) - (s-1)\rho_0(t)N(t).$$

The second term on the right-hand side of the equation contains the density of cells without ecDNA $\rho_0(t)$. We do not have a general solution for this expression, but we will see later that $\rho_0(t \to \infty) \to 0$. Consequently, for sufficiently large $N$ the tumour population will grow exponentially with $N_{s>1} = e^{st}$. Or, if we compare the relative change of fitness at any given time $t$ we get

$$\mathrm{Log}[N_{s>1}(t)] - \mathrm{Log}[N_{s=1}(t)] = (s-1)t.$$

In the initial phase of tumour growth, the term $-(s-1)\rho_0(t)N(t)$ in above equation cannot be neglected and the growth will be in the interval

$$t \leq \mathrm{Log}[N_{s>1}(t)] \leq st$$

slowly approaching the slope of $st$ with increasing time.

### 3.2 Dynamics of Moments of ecDNA copies under positive selection

In the following we discuss the dynamics of Moments for ecDNA under positive selection. Following the steps above and using the generalised Master equation for the selection case, we find the following dynamic equation for the Moments

$$\frac{\mathrm{d}M^{(l)}(t)}{\mathrm{d}t} = s\left.\frac{\mathrm{d}M^{(l)}(t)}{\mathrm{d}t}\right|_{s=1} + (s-1)\rho_0 M^{(l)}(t).$$

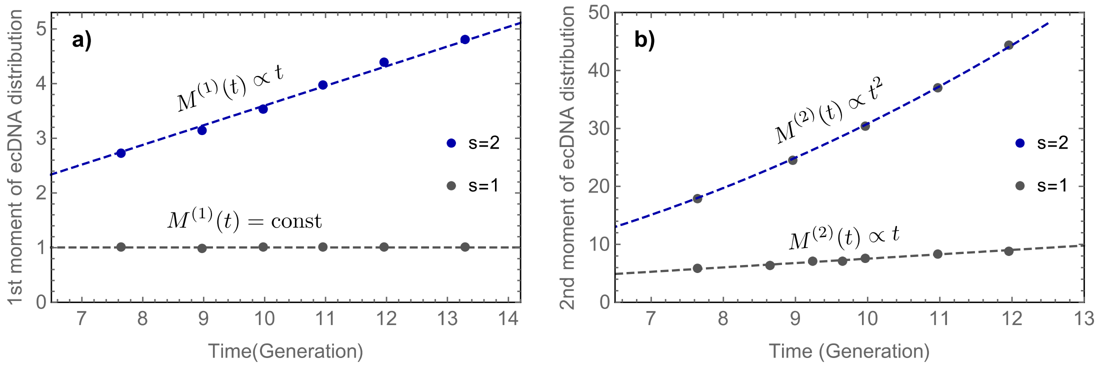


**Figure SI 8.** **a)** First and **b)** second moment of the ecDNA copy number distribution. In the neutral case ($s = 1$, grey) the mean number of ecDNA copies remains constant and the variance increases linearly in time. Under positive selection ($s = 2$, blue) the mean number of ecDNA copies increases in time. Stochastic simulations (points) are in very good agreement to theoretical predictions of polynomial increasing moments with time (dashed lines).

This implies for the first moment $\frac{\mathrm{d}M^{(1)}(t)}{\mathrm{d}t} = (s-1)\rho_0 M^{(1)}(t)$, which then can be solved for the first moment

$$M^{(1)}(t) = e^{(s-1)\int_0^t d\tau \rho_0(\tau)}.$$

Importantly, for positive selection we have $s > 1$ and therefore $s - 1 > 0$. Furthermore, the integral is strictly positive, such that the first moment is expected to increase over time. In other words, in a growing tumour population with ecDNA under positive selection, we expect the average ecDNA copy number per cell to increase in time. This is in contrast to the neutral case, where the average ecDNA copy number is expected to remain constant over time.

Similarly, the dynamic equation for the second moment becomes $\frac{\mathrm{d}M^{(2)}(t)}{\mathrm{d}t} = M^{(1)}(t) + (s-1)\rho_0 M^{(2)}(t)$ and we find

$$M^{(2)}(t) = tM^{(1)}(t).$$

The second moment is increasing as well, but now with an additional factor $t$ compared to the neutral case. Similar to the argument above, it follows that higher moments follow the form

$$M^{(l)}(t) = P_l(t) e^{(s-1)\int_0^t d\tau \rho_0(\tau)} \sim t^{l-1} M^{(1)}(t).$$

### 4.1 Deterministic population dynamics

We have in the chapters above discussed stochastic aspects of the ecDNA copy number distribution for positive and neutral selection. Another question of interest is how the fraction of cells with and without ecDNA change in a growing tumour population. We therefore change the formulation of our mathematical model to a more coarse-grained picture and only consider cells with ecDNA $N^+(t)$ and cells without ecDNA $N^-(t)$. For cells with ecDNA, we do not distinguish between different copy number states. With the notation of the former chapters, we identify $N^-(t) = N_0(t)$ and $N^+(t) = \sum_{k=1}^{\infty} N_k(t)$.

We can write for the change of these cells in time $t$

$$\frac{\mathrm{d}N^-(t)}{\mathrm{d}t} = N^-(t) + v\big(N^+(t)\big)N^+(t)$$
$$\frac{\mathrm{d}N^+(t)}{\mathrm{d}t} = N^+(t) - v(N^+(t))N^+(t)$$

where $v(N^+(t))$ is the rate at which cells with ecDNA lose all ecDNA copies by chance due to complete asymmetric random ecDNA segregation (one daughter cell inherits all copies of ecDNA, while the other cell does not inherit any). Looking at the fraction of cells with ecDNA $f^-(t) = \frac{N^-(t)}{N^+(t)+N^-(t)}$, we can write

$$\frac{\mathrm{d}}{\mathrm{d}t}\left(\frac{N^-(t)}{N^+(t)+N^-(t)}\right) = \frac{\mathrm{d}}{\mathrm{d}t} f^-(t) = (1 - f^-(t))\, v\big(N^+(t)\big)$$

Rearranging terms gives

$$v\big(N^+(t)\big) = 1 - \frac{1}{N^+(t)}\frac{\mathrm{d}N^+(t)}{\mathrm{d}t}$$

and thus we can find for the fraction of cells without ecDNA the following relation

$$\frac{1}{1 - f^-(t)}\frac{\mathrm{d}f^-(t)}{\mathrm{d}t} + \frac{1}{N^+(t)}\frac{\mathrm{d}N^+(t)}{\mathrm{d}t} = 0.$$

This equation can be integrated by separation of variables. With the initial condition $N^+(0) = 1$ and $f^-(0) = 0$ the number of cells with ecDNA is given by

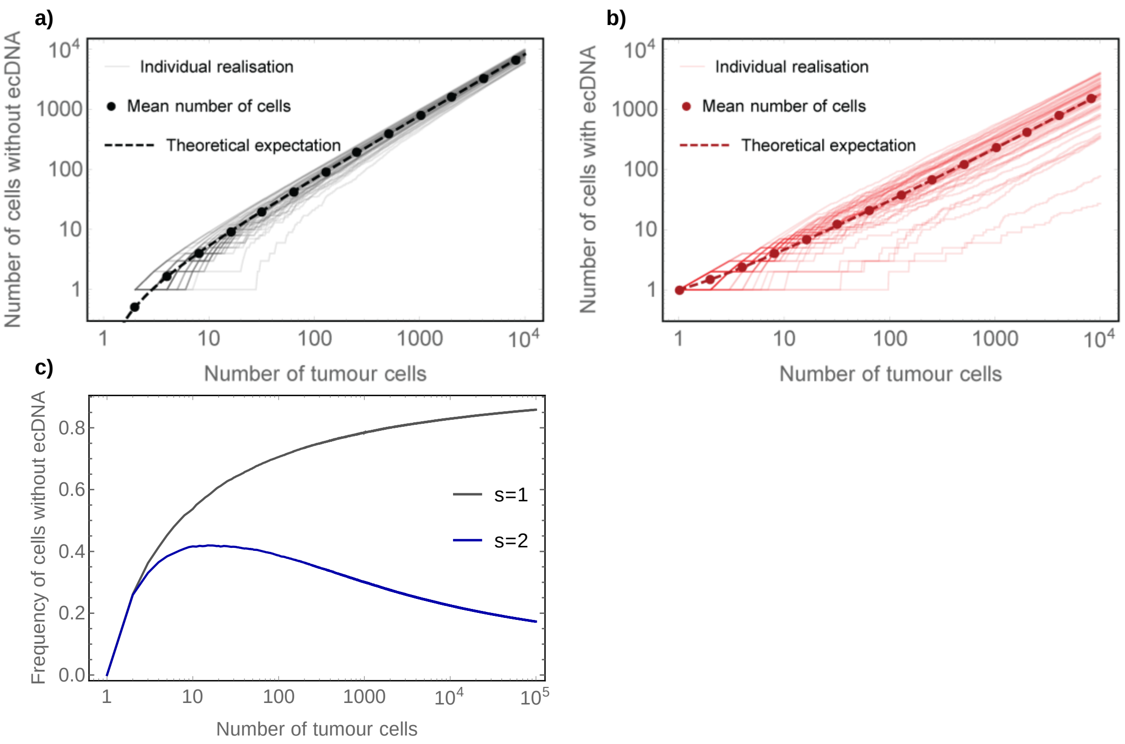


**Figure SI 9.** Comparison of average deterministic dynamics of cells **a)** without and **b)** with copies of ecDNA for neutral ecDNA dynamics ($s = 1$). Dots show the average dynamics of neutral stochastic simulations, lines are individual realisation of the same neutral stochastic process and dashed lines show analytical predictions. Between tumour variation is considerable, especially for small tumour populations. **c)** Fraction of cells without ecDNA over time. In the neutral case $s = 1$ the tumour will be dominated by cells without ecDNA, also the fitness of cells with and without ecDNA is the same. Under strong positive selection, where cells with ecDNA have a selection advantage $s = 2$, the frequency of cells without ecDNA approaches 0. Even for strong positive selection we observe a transient increase of cells without ecDNA.

$$N^+(t) = \big(1 - f^-(t)\big)e^t.$$

Stochastic simulations show that for neutral dynamics, $\frac{N^-(t)}{N^+(t)} = \frac{1}{2}t$ and therefore the fraction of cells without ecDNA changes according to

$$f^-(t) = \frac{N^-(t)}{N^+(t) + N^-(t)} = \frac{1}{\frac{N^+(t)}{N^-(t)} + 1} = \frac{1}{\frac{2}{t} + 1} = \frac{t}{2 + t}.$$

We see that $f^-(0) = 0$ and $f^-(t \to \infty) \to 1$, in the long run a growing population with neutral ecDNA elements will be dominated by cells without ecDNA. This can also be seen from the fraction of cells carrying ecDNA. From the simple condition $f^-(t) + f^+(t) = 1$ we find

$$f^+(t) = 1 - \frac{t}{2 + t} = \frac{2}{2 + t} = \frac{2}{2 + \mathrm{Log}[N]}.$$

Also, the number of cells with ecDNA continuously decreases in the neutral case, the decrease is proportional to $\sim\text{Log}^{-1}[N]$ and thus relatively slow. For example, in a population of $10^3$ cells, the expected fraction would be $22\%$, in a population of $10^6$ cells the fraction becomes 13% and in a population of $10^{11}$ cells it is 7%. With single cell resolution, we might expect to detect low levels of neutral ecDNA copies in tumour populations.

The population dynamics changes when ecDNA is under positive selection. As previously, we introduce a selection coefficient $s > 0$, with $s = 1$ corresponding to neutral selection and $s > 1$ to a selective advantage of cells carrying ecDNA. The population level dynamics now changes to

$$\frac{\mathrm{d}N^-(t)}{\mathrm{d}t} = N^-(t) + sv\big(N^+(t)\big)N^+(t)$$

$$\frac{\mathrm{d}N^+(t)}{\mathrm{d}t} = sN^+(t) - sv(N^+(t))N^+(t)$$

Following the same steps as above, this can be transformed in a single set of equations

$$(s-1)f^-(t) + \frac{1}{1-f^-(t)}\frac{\mathrm{d}f^-(t)}{\mathrm{d}t} + \frac{1}{N^+(t)}\frac{\mathrm{d}N^+(t)}{\mathrm{d}t} = s$$

Again, this equation can be formally integrated by the separation of variables and we get

$$N^+(t) = (1 - f^-(t))e^{st-(1-s)\int_0^t f^-(\tau)\mathrm{d}\tau}$$

A closed solution is more challenging in the selection case as we do not have a closed expression for $\int_0^t f^-(\tau)\mathrm{d}\tau$. However, we find numerically $f^-(t \to \infty) \to 0$ and thus for sufficiently long time, the number of cells with ecDNA grows with $N^+(t) \approx e^{st}$. A tumour population with ecDNA copies under positive selection, will be dominated by cells carrying ecDNA.

#### 4.2 Dynamic predictions of ecDNA under neutral vs positive selection

In the previous chapter, we have discussed the stochastic dynamics of the ecDNA copy number distribution as well as the deterministic aspect of the population dynamics of cells with and without ecDNA in exponentially growing populations. This leads to three major predictions that differ between cell populations under neutral dynamics or positive selection.

(i) Fraction of cells with and without ecDNA: Theory predicts that the fraction of cells with ecDNA approaches 0 under neutral dynamics and approaches 1 if ecDNA is under positive selection. The rate of convergence depends on the strength of selection. In all patient and cell line samples, we find a very high fraction of cells with ecDNA, suggesting positive selection.

(ii) Average ecDNA copy number per cell: Theory predicts that the average ecDNA copy number per cell increases in time, if ecDNA is under positive selection and remains on average at 1 if ecDNA is under neutral selection. In all patient and cell line samples we find average ecDNA copy numbers $\gg 1$, suggesting positive selection.

(iii) Power law scaling of the ecDNA copy number distribution: Theory predicts a 1/copy number scaling of the ecDNA copy number distribution for both neutral and positive selection. We find this scaling in patient and cell line experiments. However, the ecDNA copy number distribution shifts towards higher copy number under positive selection and consequently, the power law tail is shifted towards higher ecDNA copy number as well. We observe these behaviours in patient and cell line experiments.

.
